# Comprehensive bioinformatic analysis of newly sequenced *Turdoides affinis* mitogenome reveals the persistence of translational efficiency and dominance of NADH dehydrogenase complex-I in electron transport system over Leiothrichidae family

**DOI:** 10.1101/2020.01.23.917716

**Authors:** Indrani Sarkar, Prateek Dey, Sanjeev Kumar Sharma, Swapna Devi Ray, Ram Pratap Singh

**Affiliations:** National Avian Forensic Laboratory, Sálim Ali Centre for Ornithology and Natural History, Anaikatty, Coimbatore – 641108, Tamil Nadu, India

**Author notes:** Corresponding Author: Dr. Ram Pratap Singh, Senior Scientist, Sálim Ali Centre for Ornithology and Natural History, Anaikatty, Coimbatore – 641 108, Tamil Nadu, India.

**Keywords:** Phylogeny, mitochondrial DNA, birds, tRNA, Control region

## Abstract

Mitochondrial genome provides useful information about species with respect to its evolution and phylogenetics. We have taken the advantage of high throughput next-generation sequencing technique to sequence the complete mitogenome of Yellow-billed babbler (*Turdoides affinis*), a species endemic to Peninsular India and Sri Lanka. Both, reference-based and *de-novo* assemblies of mitogenome were performed and observed that *de-novo* assembled mitogenome was most appropriate. The complete mitogenome of yellow-billed babbler (assembled *de-novo*) was 17,671 bp in length with 53.2% AT composition. Thirteen protein-coding genes along with 2 rRNAs and 22 tRNAs were detected along with duplicated control regions. The arrangement pattern of these genes was found conserved among Leiothrichidae family mitogenomes. Downstream bioinformatics analysis revealed the effect of translational efficiency and purifying selection pressure over all the thirteen protein-coding genes in yellow-billed babbler mitogenome. Moreover, genetic distance and variation analysis indicated the dominance of NADH dehydrogenase complex-I in the electron transport system of *T. affinis*. Evolutionary analysis revealed the conserved nature of all the protein-coding genes across Leiothrichidae family mitogenomes. Our limited phylogenetics results suggest that *T. affinis* is closer to *Garrulax*.

## Introduction

Aves are one of the most diverse vertebrate classes with a huge number of species having a broad range of ecological behavior and complex morphology, all of which make it difficult to solve the riddles regarding their taxonomy along with phylogenetic and evolutionary relationship^1–3^. New and advanced scientific techniques have emerged to solve these riddles. For the last few years, genome sequencing has become more popular to obtain huge information on evolutionary history and revising the clustering pattern of traditional taxonomy^4^. Mitochondrial DNA with some of its inherent properties like small genome size, absence of extensive recombination frequency, simple structure of genome, maternal inheritance along with rapid evolutionary rate are now extensively utilized in taxonomic and phylogenetic studies of vertebrates^5–10^. Furthermore, it has been reported that, complete mitogenomes retain more information than a single gene regarding the evolutionary history of the taxon and also provide consistent results compared to nuclear genes^11^. This also reduces the effect of homoplasy and frequent stochastic errors in phylogenetic studies^11^.

Yellow-billed bababler (*Turdoides affinis*) is one of the most common birds in India^12^. They are distributed in the southern peninsular India including the southern part of Maharashtra, Chattisgarh and Andhra Pradesh^12^. The taxonomic classification of this bird is quite dubious. Previously all babbler and allies were considered under Timaliidae family^13^. Recent classification studies have split this family into five discrete families^13^. Three of them, Leiothrichidae, Pellorneidae and Timaliidae consisted of traditional babblers while Zosteropidae included mainly *Yuhina* along with some other minor species and Sylviidae grouped all the *Sylvia warblers*^14^. Among these five distinct families, Leiothrichidae represented the largest group consisting of 125 species distributed mainly in the Sino-Himalayan and South-Eastern parts of Asia^14^. A study on the Leiothrichidae family suggested their origin prior to the Miocene–Pliocene boundary, which is well known for its noteworthy climatic turmoil in Asia^13^.Hence, it is imperative to conduct in-depth molecular studies on this family to know some very appealing unknown facts regarding their taxonomical and phylogenetic relationships.

This Leiothrichidae family mainly consisted of *Grammatoptila*, *Garrulax*, *Trochalopteron*, *Turdoides* and *Argya*^14^. Polyphyletic nature was observed among members of *Turdoides* due to which further taxonomic classification was done and some species were resurrected from this genus^14–16^. Yellow-billed babbler which is considered to be under *Turdoides* has been recently proposed to be under *Argya* based on revised taxonomy of Leiothrichidae^14^; however, more investigations are required to confirm this. Yellow-billed babbler generally live in flocks of seven to ten members continuously squeaking, chattering and chirping. Helpers are seen generally assisting parents in nest building, chick feeding and maintaining nest sanitation as a cooperative breeding character^17–19^. Interestingly, the close relatives of *T. affinis* for instance, *Garrulax*, *Leiothrix*, *Liocichla*, *Minla* and *Trochalopteron* have still not been reported to show the cooperative breeding behaviour indicating a divergence of *T. affinis* at least from the behavioural ecology perspective^18^.

Untill now, the complete mitogenomes of various species (including *Garrulax affinis*^20^, *G. albogularis*, *G. canorus*, *G. canorus* voucher AHNU, *G. cineraceus*^21^, *G. elliotii*^22^, *G. formosus*^23^, *G. ocellatus*^24^, *G. perspicillatus*^25^, *G. poecilorhynchus* voucher B33^26^, *G. sannio*^27^, *G. sannio* voucher GSAN20150704V3, *Trochalopter onmilnei, Leiothrix argentauris*, *L. lutea*, *Liocichla omeiensis*^28^ and *Minla ignotincta*^29^) from Leiothrichidae family have been reported. However, complete mitogenome of birds representing the genus *Turdoides* is yet to be revealed.

In this study, we have sequenced and described the complete mitochondrial genome of *T.affinis* obtained using two approaches such as reference-based assembly and *de-novo* assembly^30–31^. We employed two different genome assembly approaches to align the same mitogenome to quantify the differences occurring due to the alignment approach and their effect on mitogenomic parameters. In addition, we have performed a detailed comparative analysis on available mitogenomes of the family Leiothrichidae to understand the overall species-specific differences including some potent parameters such as codon usage and evolutionary history.

## Materials and Methods

### Sample collection and DNA extraction

A fresh road-killed specimen of *T. affinis* was collected from Anaikatty Hills in Coimbatore district of Tamil Nadu, India (Fig. 1), and transported immediately to the lab. Prior permission for roadkill collection was obtained from Tamil Nadu Forest Department (Ref.No.WL5 (A)/2219/2018; Permit No. 14/2018). Muscle tissue was sampled from the specimen and stored in DESS buffer (20% DMSO, 0.25M tetra-sodium EDTA, Sodium Chloride till saturation, pH 7.5) at −20°C for further processes. About 50 milligram of the muscle tissue in DESS buffer was taken and added to 500 µl of lysis buffer (10 mM Tris-pH 8.0, 10 mM EDTA-pH 8.0, 100 mM NaCl) and homogenized thoroughly. To the homogenate,80 μl of 10% SDS and along with 20 μl of Proteinase K (20 mg/ml) was added and incubated overnight at 55°C. The DNA extraction was performed the following day using suitable volumes of Phenol, Chloroform and Isoamyl alcohol. The DNA pellet obtained was suspended in 100 µl of Tris-EDTA buffer (Sigma-Aldrich, USA) and quantified using spectrophotometer (DeNovix, USA) and Qubit 4 Fluorometer (ThermoFisher Scientific, USA). The quality of the DNA extracted was assessed by running it on 1% agarose gel stained with ethidium bromide intercalating dye.

**Figure 1.**
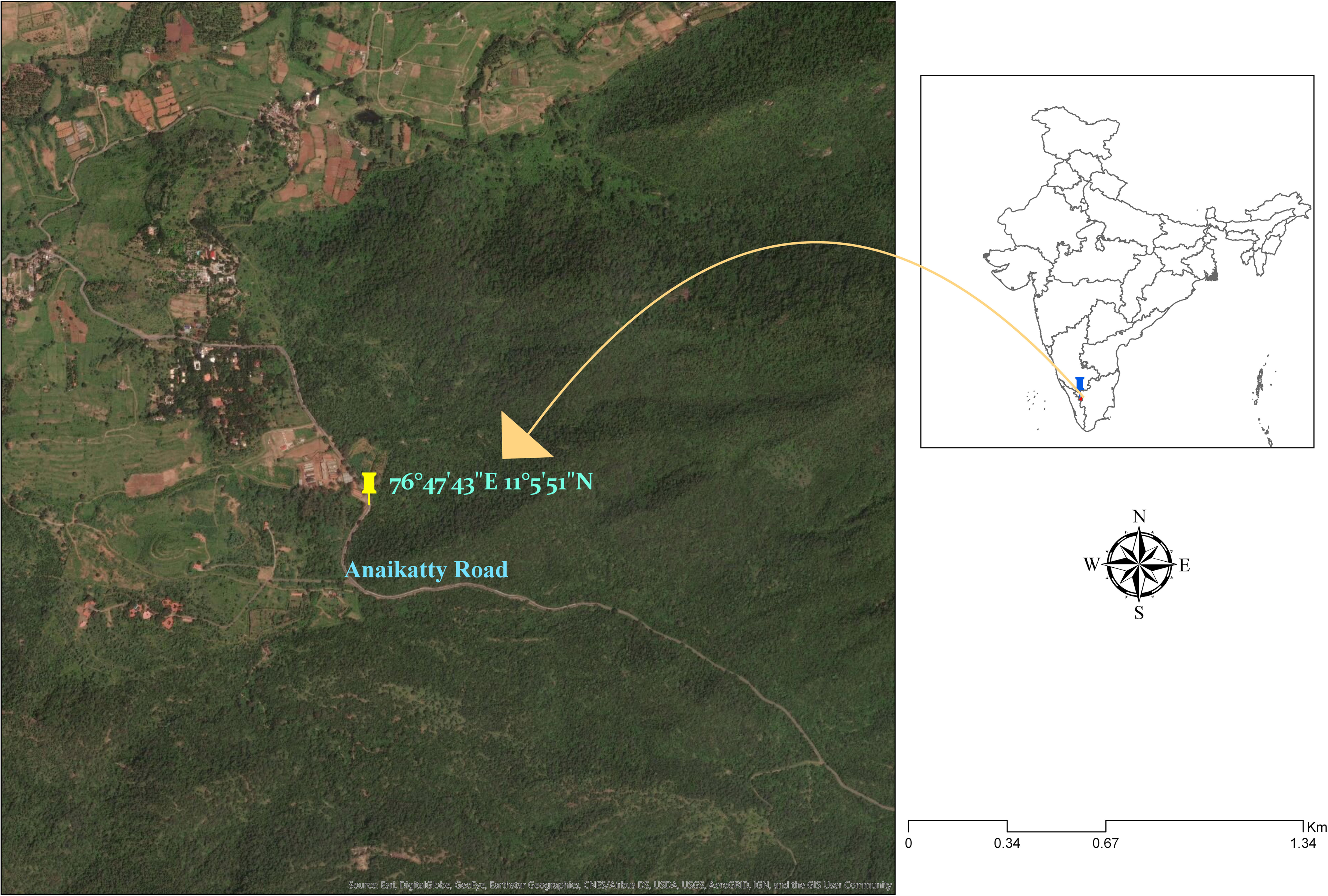
Map showing sample collection area.

### Library preparation

For library preparation, 700 nanograms of extracted DNA was utilized as starting material in NEBNext Ultra II DNA Library Prep kit for Illumina (New England Biolabs, USA). The DNA was fragmented using focused ultrasonicator (Covaris M220, USA) until the desired length of 270-300 base pairs was obtained. The fragmented DNA size was analyzed by running it in Fragment Analyzer (Agilent, USA) making sure that the size of the majority of DNA fragments is between 270-300 base pairs. Adaptor ligation was then carried out in a thermocycler following the “NEBNext Ultra II DNA Library Prep kit for Illumina” protocol using dual indexed primers present in the kit for creation of pair end libraries. After the ligation reaction was completed, the size selection of Adaptor-ligated DNA was carried out using NEBNext Sample Purification Beads (New England Biolabs, USA) followed by PCR enrichment of Adaptor-ligated DNA following the manufacturer’s protocol. After the clean-up of the enriched DNA, it was again analysed for the required concentration and mean peak size in a Fragment Analyzer (Agilent, USA). The enriched DNA library fragments were subjected to sequencing in Illumina NextSeq 550 (Illumina, Inc., USA) using Illumina High Output Kit for NextSeq 500/550 (Illumina, Inc., USA). A PhiX control library (Illumina, Inc., USA) was also subjected to sequencing along with the sample DNA library as an internal control. At the end of the sequencing run, high quality paired end reads were obtained, and further bioinformatics analysis was performed.

### Assembly and annotation of the mitochondrial genome

IlluminaNextSeq 550 produced 88,52,137 raw reads from whole-genome library. Cutadapt tool^32^ was used to trim the adapter and lowquality reads with a Phred (Q) score of 30 was selected for further analysis. Finally we got 12,97,736 high quality reads after down sampling with Seqtk (https://github.com/lh3/seqtk) which were used for assembly. *De novo* assembly was performed using SPAdes-3.11.1 software with default parameters. MITOS online server (http://mitos.bioinf.uni-leipzig.de/index.py) was used for annotation of the mitogenome. Reference-based assembly was also performed as described in supplementary file 1.

### Phylogenetic analysis

Phylogenetic analysis was performed on the available whole mitogenomes of various species of the Leiothrichidae family. We prepared two complete mitogenome based phylogenetic trees, one with *T. affinis* mitogenome assembled through reference-based assembly and another with *de-novo* assembly using Maximum likelihood algorithm with 1000 bootstrap values in ClustalW implemented with Mega ver. 7.

### Sequence analysis of mitogenome

The complete mitogenome of *T. affinis* was compared with other Leiothrichidae family avian species whose complete mitochondrial genomes are available at NCBI including *Garrulax affinis*^20^, *G. albogularis*, *G. canorus*, *G. canorus* voucher AHNU, *G. cineraceus*^21^, *G. elliotii*^22^, *G. formosus*^23^, *G. ocellatus*^24^, *G. perspicillatus*^25^, *G. poecilorhynchus* voucher B33^26^, *G. sannio*^27^, *G. sannio* voucher GSAN20150704V3, *Trochalopteron milnei, Leiothrix argentauris*, *L. lutea*, *Liocichla omeiensis*^28^ and *Minla ignotincta*^29^. These sequences were downloaded from NCBI website and used for further comparative analysis. The protein-coding genes along with tRNAs and rRNAs were aligned to examine whether any rearrangements persist among these mitogenomes. The initial and terminating codons of all the protein-coding genes were curated through NCBI ORF finder (https://www.ncbi.nlm.nih.gov/orffinder/). Circular genome views were obtained by CG view server^33^. The boundary of the control region (CR) was determined^34^. Detailed codon usage analysis of the select mitogenomes was performed using CodonWsoftware^35^. The studied codon usage parameters include Relative Synonymous codon Usage (RSCU), Effective number of Codons (ENc) and frequency of G+C at the third position of codons (GC3s). Different codon composition indices of individual genes, for example total GC content as well as the frequency of each nucleotide at the third position of codons (A3, T3, G3 and C3) were also estimated. R-based heatmaps were generated based on the overall codon usage and amino acid usage analysis. Skew values for AT[(A-T)/(A+T)] and GC [(G-C)/(G+C)] were calculated^36^ using DAMBE software (http://dambe.bio.uottawa.ca/DAMBE/dambe.aspx). Tandem Repeats Finder software (https://tandem.bu.edu/trf/trf.html) with its default settings was employed to detect any tandem repeats within the mitogenomes. A BlastN based approach to find intra-genomic duplication of large fragments or interspersed repeats was employed^37^, where each mitogenome was searched against itself with an e-value of 1e-10. This analysis detected a negligible number of interspersed repeats, hence was not evaluated further.

### Estimation of translational efficiency

This parameter measures the competence of codon-anticodon interactions indicating the accuracy of the translational machinery of genes in the absence of preferred codon set information. We calculated the translational efficiency according to the following equation^38^:

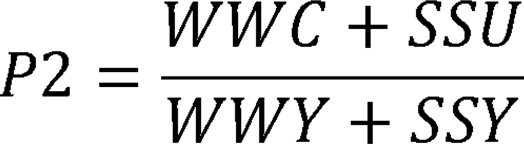

where, W= A or U, S=C or G and Y=C or U.

P2 > 0.5 indicates the existence of translational selection.

### RSCU based cluster analysis and putative optimal codons

Generally, highly expressed genes utilize a specific set of codons termed as optimal codons. Due to the preferential use of this set of codons their Enc value lowers down in contrast to lowly expressed genes, which restrain more rare codons with higher Encvalue^35^. We identified the optimal codons of all investigated species from their RSCU values. RSCU =1 indicated unbiased codon usage whereas; RSCU > 1 and RSCU < 1 indicated a higher and lower usage frequency of that particular codon respectively ^35^.

### Evolutionary analysis

The ratio (⍰) of non-synonymous substitution rate per synonymous site (ka) to synonymous substitution rate per non-synonymous site (ks) has been reported to be an excellent estimator of evolutionary selection pressure or constrain on protein-coding genes. ⍰>1 stands for positive Darwinian selection (diversifying pressure), on the contrary, ⍰<1 signifies purifying or refining selection. At neutral evolutionary state, the value of ⍰becomes 1 symbolizing the equal rate of both synonymous and non-synonymous substitution^39^. The mean genetic distance of the annotated protein-coding genes of the studied mitogenomes were calculated in terms of Kimura-2-parameter (K2P) substitution model and evolutionary rate (⍰) was calculated by DnaSPver 6.12.03 software^40^.

## Results and Discussion

### Comparison of *T. affinis* mitogenome assembled using reference-based and *de-novo* assembly approach

In this study, we performed both, reference-based assembly and *de-novo* assembly of the newly sequenced mitogenome of *T. affinis* and found a considerable difference in the results between these two approaches. In reference-based assembly, the total size of the mitogenome was 16,861 bp with 47% GC and 53% AT (Supplementary file 1) whereas the *de-novo* assembly resulted in 17,671 bp long mitogenome with 53.2% AT and 46.80% GC(Fig. 2, Table 1). AT and GC skewness were 0.13 and −0.38, respectively for *de-novo* assembly. However, reference-based assembly resulted in lower AT (0.05) and GC skew (−0.14) for *T. affinis* mitogenome. The Genbank accession number of complete *T. affinis* mitogenome (assembled *de-novo*) is MN848144.

**Figure 2.**
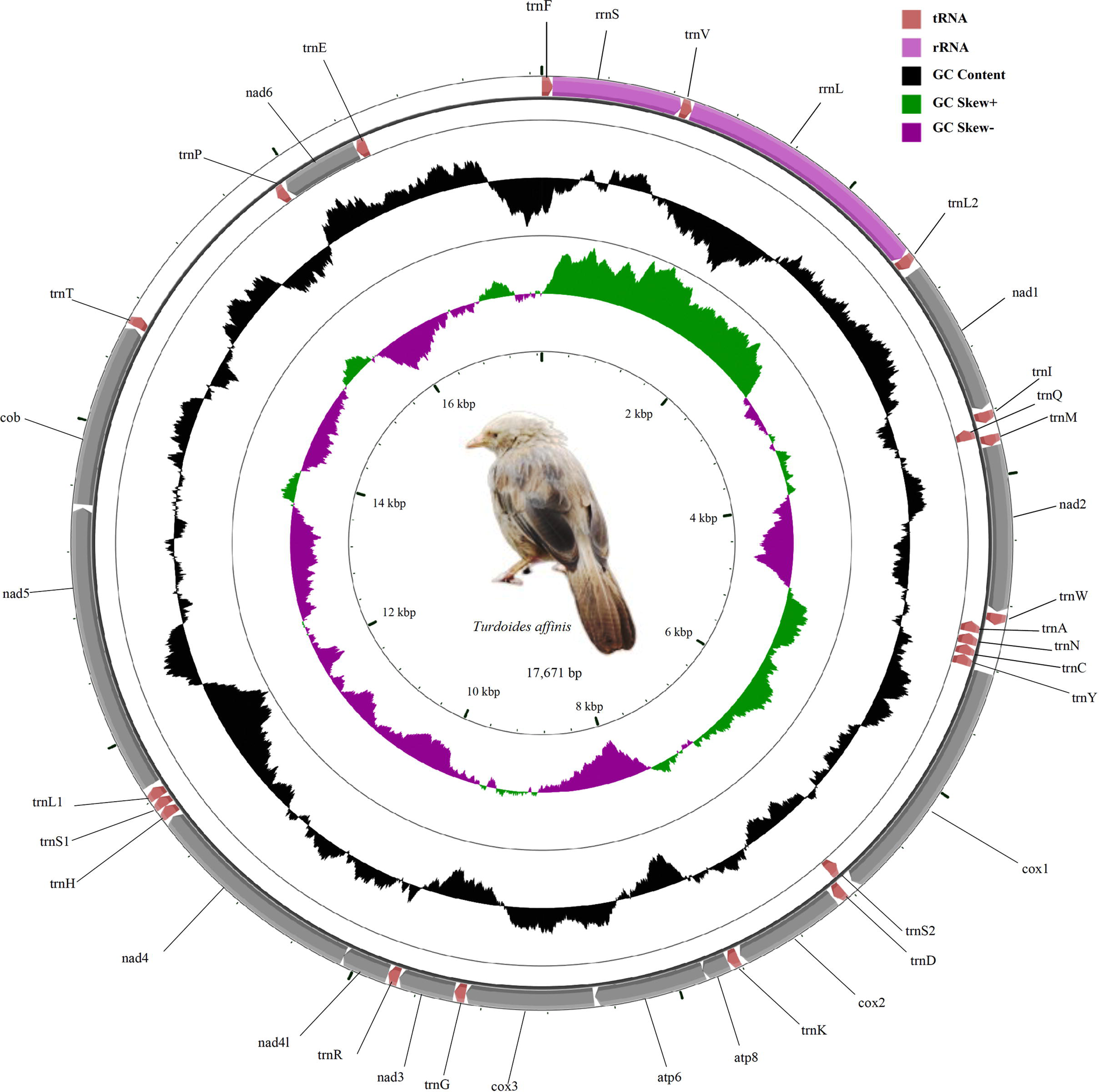
The mitochondrial genome view of *Turdoides affinis*. Gene transcription direction is indicated by arrows. Colour codes are indicated at the right upper side of the figure. tRNAs are indicated with the single letter code of amino acids. Black sliding window indicated the GC content of all the regions. GC skew has been plotted through green and violet colour sliding windows. The figure was drawn by CGView Online server (http://stothard.afns.ualberta.ca/cgview_server/) using default parameters. The photograph of *Turdoides affinis* was taken by the first author and was edited with Paint.net.

**Table 1.**
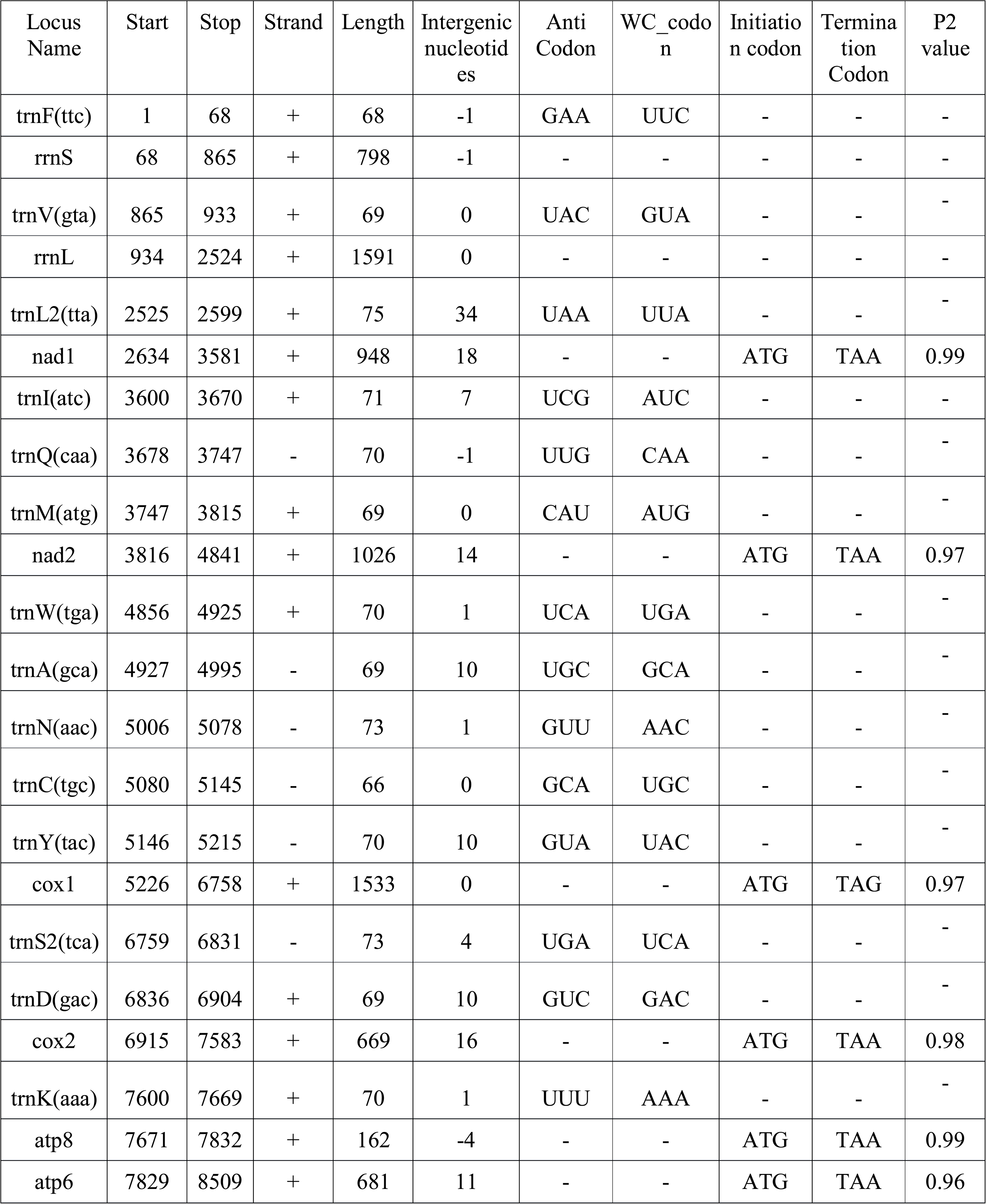
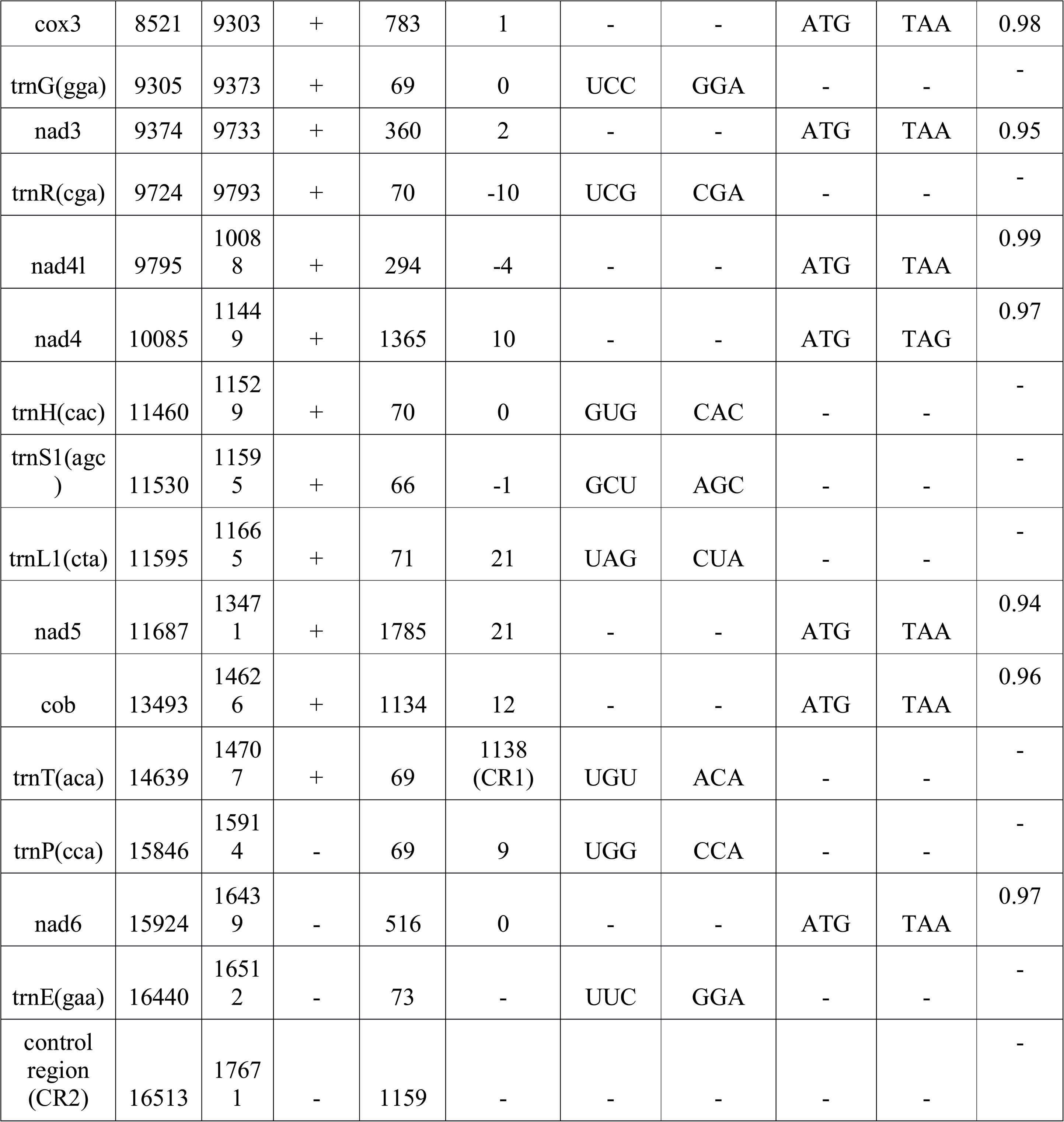
Properties of complete mitogenome of *Turdoides affinis* assembled using *de-novo* assembly approach

Two rRNA (rrnS for small subunit and rrnL for large subunit), 13 protein-coding genes (PCGs) and 22 tRNAs specified for 20 amino acids (two tRNAs each for serine and lysine) were reported in both the mitogenomes. The total length of PCGs, tRNAs and rRNAs were 11244 bp, 1539 bp and 2389 bp, respectively for *de-novo* assembly while these were 11247 bp, 1535 bp and 2578 bp respectively for reference-based assembly.

The following results were identical for both the mitogenomes. For instance, most of tRNAs (16) were distributed on the positive (+) strand except trnQ(CAA), trnA(GCA), trnN(AAC), trnC(TGC), trnY(TAC) and trnS2(TCA) that were distributed on the negative (-) strand. Both rRNAs along with all the PCGs, except nad6, were present on the negative (-) strand.

Two non-coding control regions were found and referred to as CR1 and CR2. The 5’ boundary of CR1 was trnT(ACA) and 3’ boundary was trnP(CCA) while CR2 was present between trnE(GAA) and trnF(TTC). Length of CR1 and CR2 were 1138 bp and 1159 bp, respectively in *de-novo* assembly (at parwith the other compared species) while for reference-based assembly CR1 was 825bp (less than the average CR1 length of other compared species by 300bp) and CR2 was 1539bp long (extra 390bp than the average CR2 length of compared species). The nucleotide composition of both CR1 and CR2 was calculated. AT of CR1 was 54.63% (45.37% GC) for *de-novo* assembly and 53.68% (46.32% GC) for reference-based assembly. CR2 showed 53.78% AT (46.22% GC) and 55.9% AT (44.1% GC) for *de-novo* and reference-based assembly, respectively indicating a bias towards an AT for these regions. rRNAs, tRNAs and PCGs were arranged in the following manner in both the assemblies:

trnF-rrnS-trnV-rrnL-trnL2-nad1-trnI-trnQ-trnM-nad2-trnW-trnA-trnN-trnC-trnY-cox1-trnS2-trnD-cox2-trnK-atp8-atp6-cox3-trnG-nad3-trnR-nad4l-nad4-trnH-trnS1-trnL1-nad5-cob-trnT-trnP-nad6-trnE.

These results showed considerable differences between reference-based assembly and *de-novo* assembly. Further, we performed a limited phylogenetic analysis with both the mitogenomes, and observed that *de-novo* assembled mitogenome performed better. The phylogenetic analysis of reference-based assembled mitogenome placed *T. affinis* with *Leiothrix lutea*(Supplementary file 2), whereas in case of *de-novo* assembled mitogenome based phylogeny, *T. affinis* formed a discrete group which was placed near *Garrulax affinis*(Fig. 3). The later finding is consistent with a recent report^14^ that revised the taxonomy of Leiothrichidae using a set of nuclear genes with a mitochondrial PCG as a phylogenetic marker. Though our phylogenetic results are limited because of the unavailability of complete mitogenome sequences of other groups, still it provides supporting evidences that *de novo* assembled mitogenome is more appropriate.

**Figure 3.**
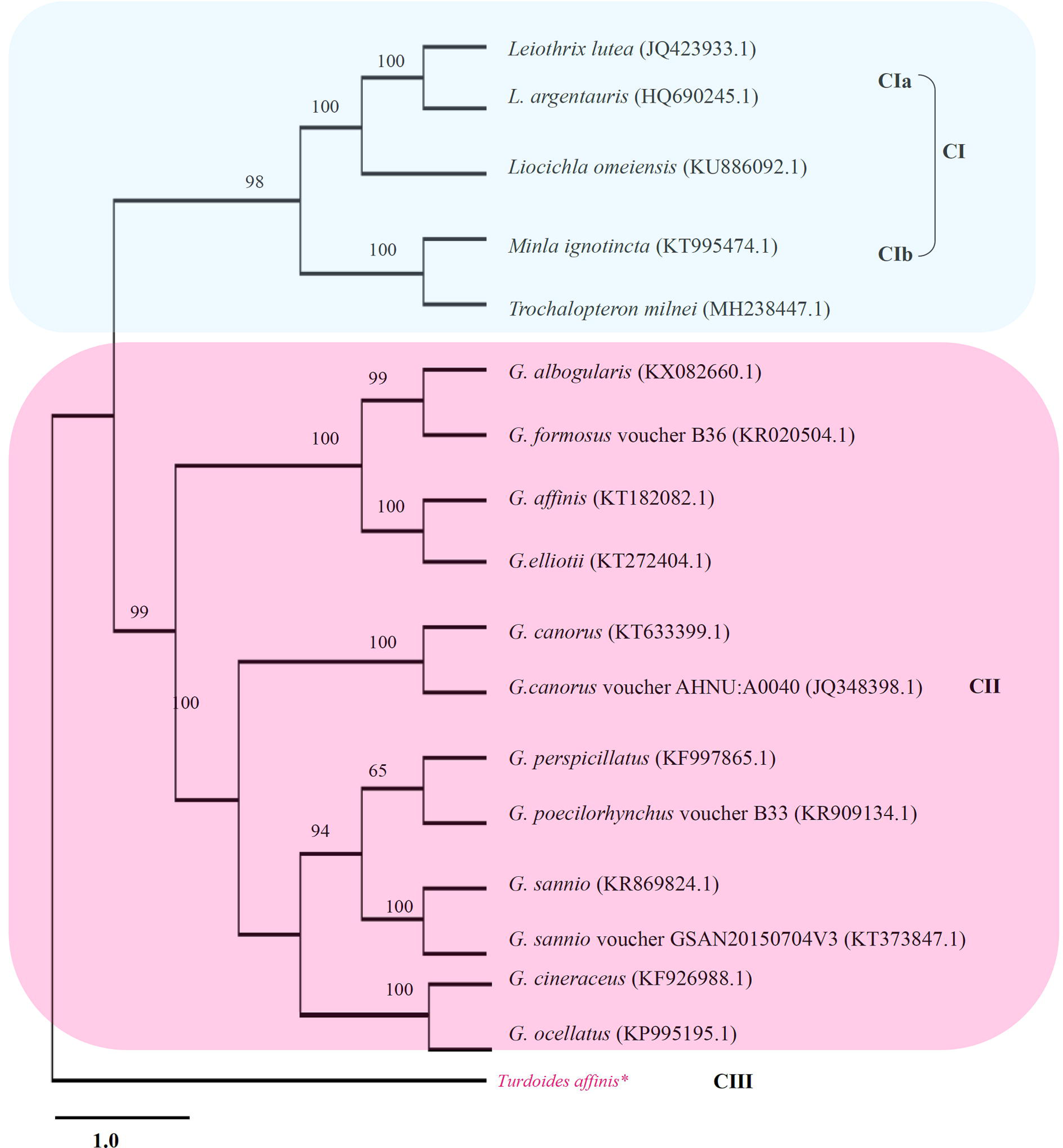
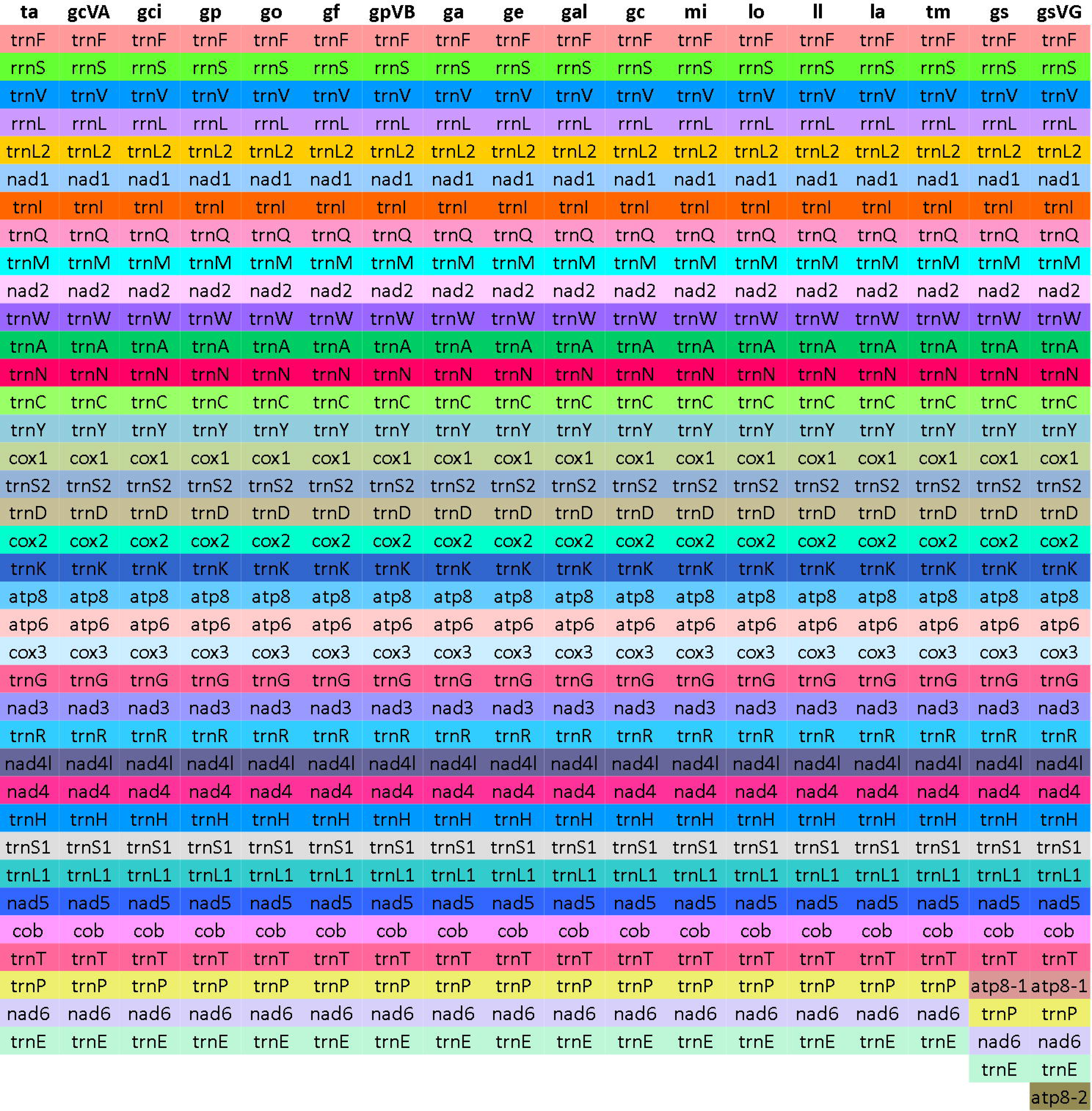
Phylogenetic tree based on complete mitogenome (*De-novo* assembled *Turdoides affinis* mitogenome). Maximum likelihood method with 1000 bootstrap value along with Kimura 2-parameter was used to generate the phylogenetic tree.

### Nucleotide composition and translational efficiency

The newly sequenced complete mitogenome of *T. affinis* was compared with other available mitogenomes from the Leiothrichidae family (Table 2). We considered *Garrulax affinis*(KT182082.1)^20^, *G. albogularis*(KX082660.1), *G. canorus*(KT633399.1), *G. canorus* voucher AHNU (JQ348398.1), *G. cineraceus*(KF926988.1)^21^, *G. elliotii*(KT272404.1)^22^, *G. formosus*(KR020504.1)^23^, *G. ocellatus*(KP995195.1)^24^, *G. perspicillatus*(KF997865.1)^25^, *G. poecilorhynchus* voucher B33 (KR909134.1)^26^, *G. sannio*(KR869824.1)^27^, *G. sannio* voucher GSAN20150704V3 (KT373847.1), *Leiothrix argentauris*(HQ690245.1), *L. lutea*(JQ423933.1), *Liocichla omeiensis*(KU886092.1)^28^, *Minla ignotincta*(KT995474.1)^29^ and *Trochalopteron milnei*(MH238447.1). Gene arrangement pattern was observed among these species and was found similar to that of *T. affinis* (Fig. 4). Values of AT skew ranged from 0.09 (*T. milnei*) to 0.13 (*Turdoides affinis*) while the GC skew ranged from −0.39 (*G. albogularis*) to −0.36 (*L. omeiensis*) (Table 2). Comparative RSCU analysis identified a set of optimal codons common among all species – GCC(A), UGC(C),UUC(F), GGA(G), AAA(K), CUA(L), UUA(L), AUA(M), CCU(P), CAA(Q), AGC(S), ACC (T) and GUA(V). Along with these GCU(A), GCA(A), GAC(D), GAA(E), CAC(H), AUU(I), CUC(L), CUU(L), AAU(N), CCC(P), CGC(R), UCU(S), ACU(T), GUC(V), UGA(W) and UAC(Y) were also frequently used in *T. affinis*. Heatmaps based on codon and amino acid usage (Fig. 5) analysis of compared mitochondrial genomes validated the aforementioned codon preference.

**Figure 4.**
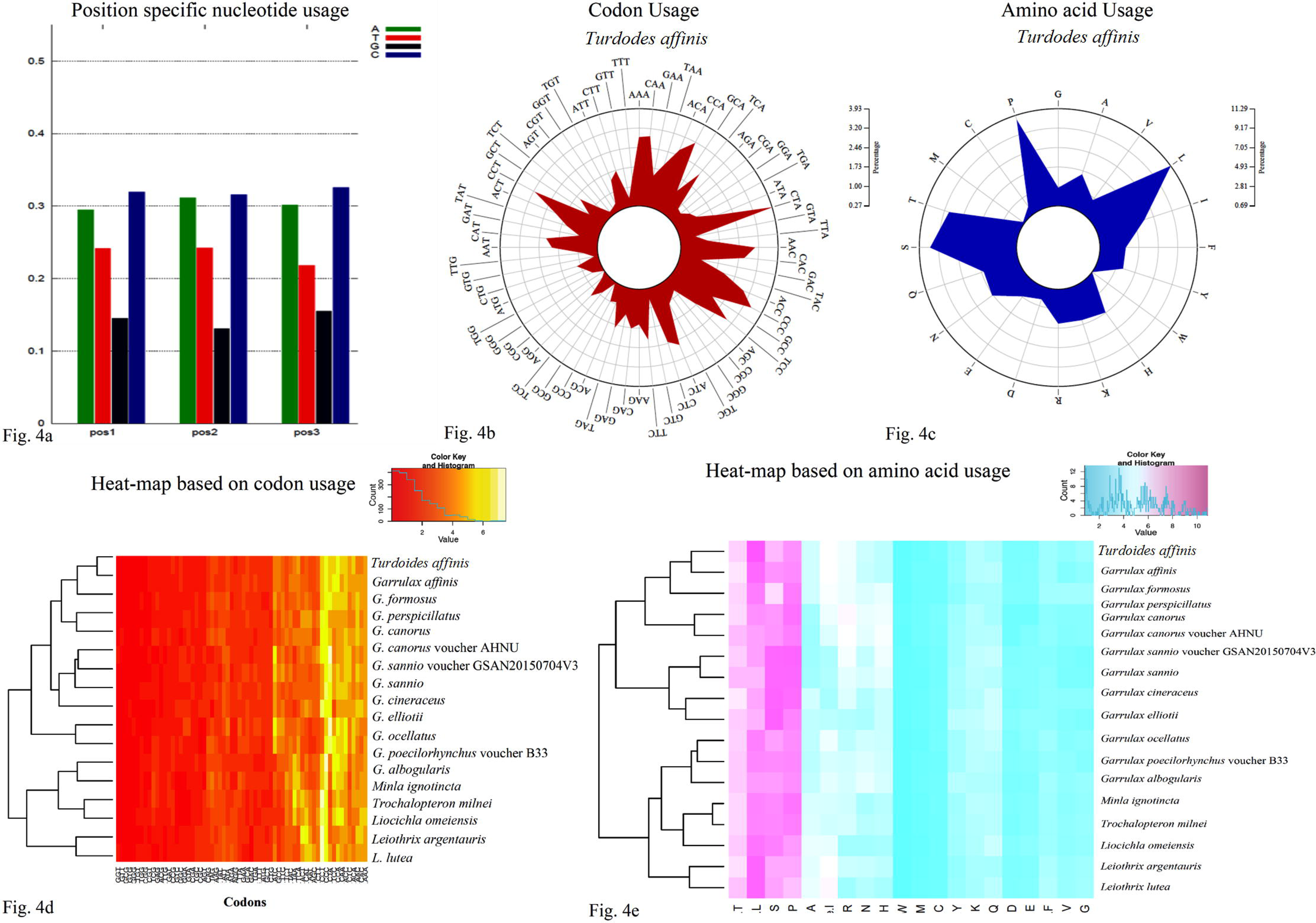
Comparison of gene orders among select complete mitogenomes of Leiothrichidae family. Organisms names are abbreviated as:ta -*Turdoides affinis*, ga – *Garrulax affinis* (KT182082.1), gal *-Garrulax albo*gularis (KX082660.1), gcVA *-Garrulax canorus* voucher AHNU:A0040 (JQ348398.1), gc – *Garrulax canorus* (KT633399.1), gci – *Garrulax cineraceus* (KF926988.1), ge – *Garrulax elliotii* (KT272404.1), gf – *Garrulax formosus* voucher B36 (KR020504.1), go – *Garrulax ocellatus* (KP995195.1), gp – *Garrulax perspicillatus* (KF997865.1), gpVB – *Garrulax poecilorhynchus* voucher B33 (KR909134.1), gsVG – *Garrulax sannio* voucher GSAN20150704V3 (KT373847.1), gs – *Garrulax sannio* (KR869824.1), la – *Leiothrix argentauris* (HQ690245.1), ll – *Leiothrix lutea* (JQ423933.1), lo – *Liocichla omeiensis* (KU886092.1), mi – *Minla ignotincta* (KT995474.1), tm – *Trochalopteron milnei* (MH238447.1). Genes were found arranged in an identical fashion except for gs and gsVG.

**Figure 5.**
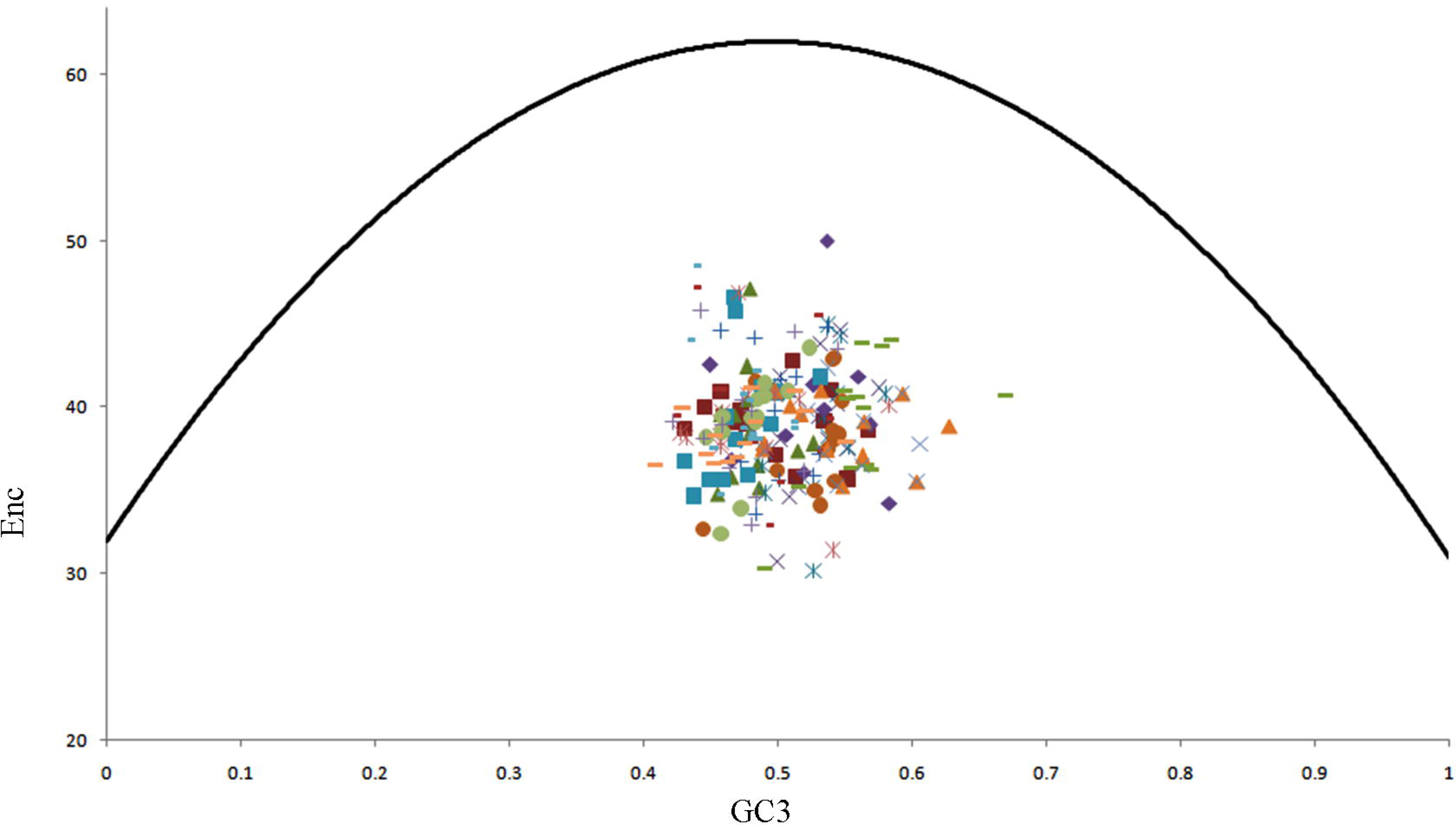
Nucleotide composition of *Turdoides affinis* mitogenome. (a) Position-specific nucleotide usage in *Turdoides affinis* mitogenome. This result was validated by the (b) roseplot based on codon usage of *Turdoides affinis* mitogenome. (c) roseplot based on amino acid usage of *Turdoides affinis* mitogenome. Heatmaps based on (d) codon usage and (e) amino acid usage.

**Table 2.**
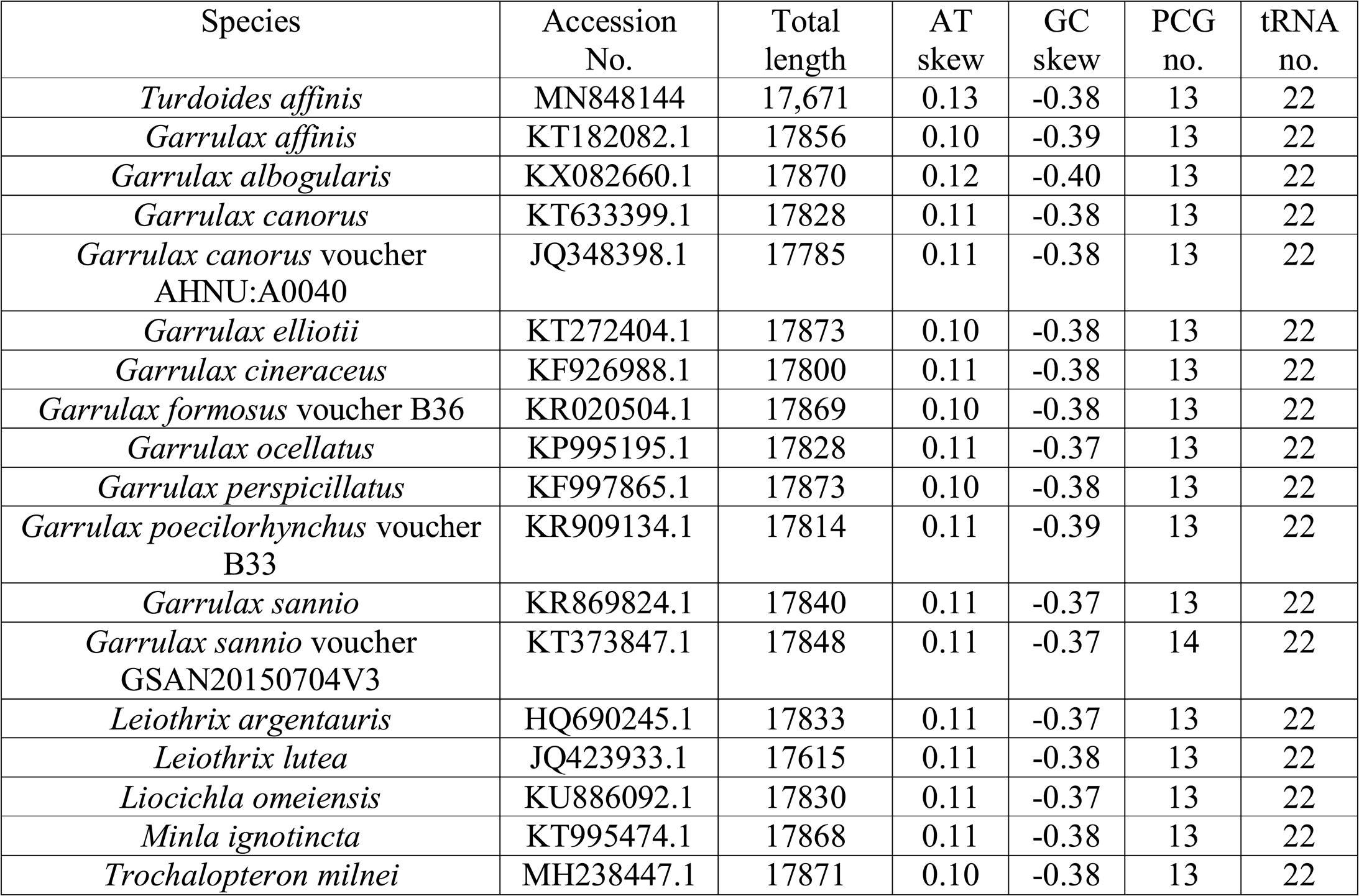
List of complete mitochondrial genomes used for comparative mitogenomics study.

GC3 vs. Enc plot analysis has been proved very efficient in predicting whether translational selection or mutational pressure persist over the genes of interest^35^. The GC3-Enc plots(Fig.6a) of protein-coding genes of the compared mitogenomes were placed well below the curve indicating the predominance of selection pressure over mutational bias. RSCU analysis revealed a higher degree of concord among Leiothrichidae mitogenomes from the codon usage perspective (Fig. 6b, Supplementary file 3). To substantiate the factors governing this codon practice GC3 vs. Enc plot analysis was done. It has been proposed that, GC3-Enc plot of genes should be placed on or above the continuous Enc curve when only mutational pressure prevails. However, in the presence of translational selection, the plots should fall well below the aforementioned curve^35^. Here the GC3-Enc plots of protein-coding genes of all studied mitogenomes were below the curve designating the influence of translational selection over those genes. Hence, we conclude that, the codon usage pattern of *T. affinis* mitogenome along with other examined species is affected by the pervasiveness of translational selection over mutational pressure.

**Figure 6.**
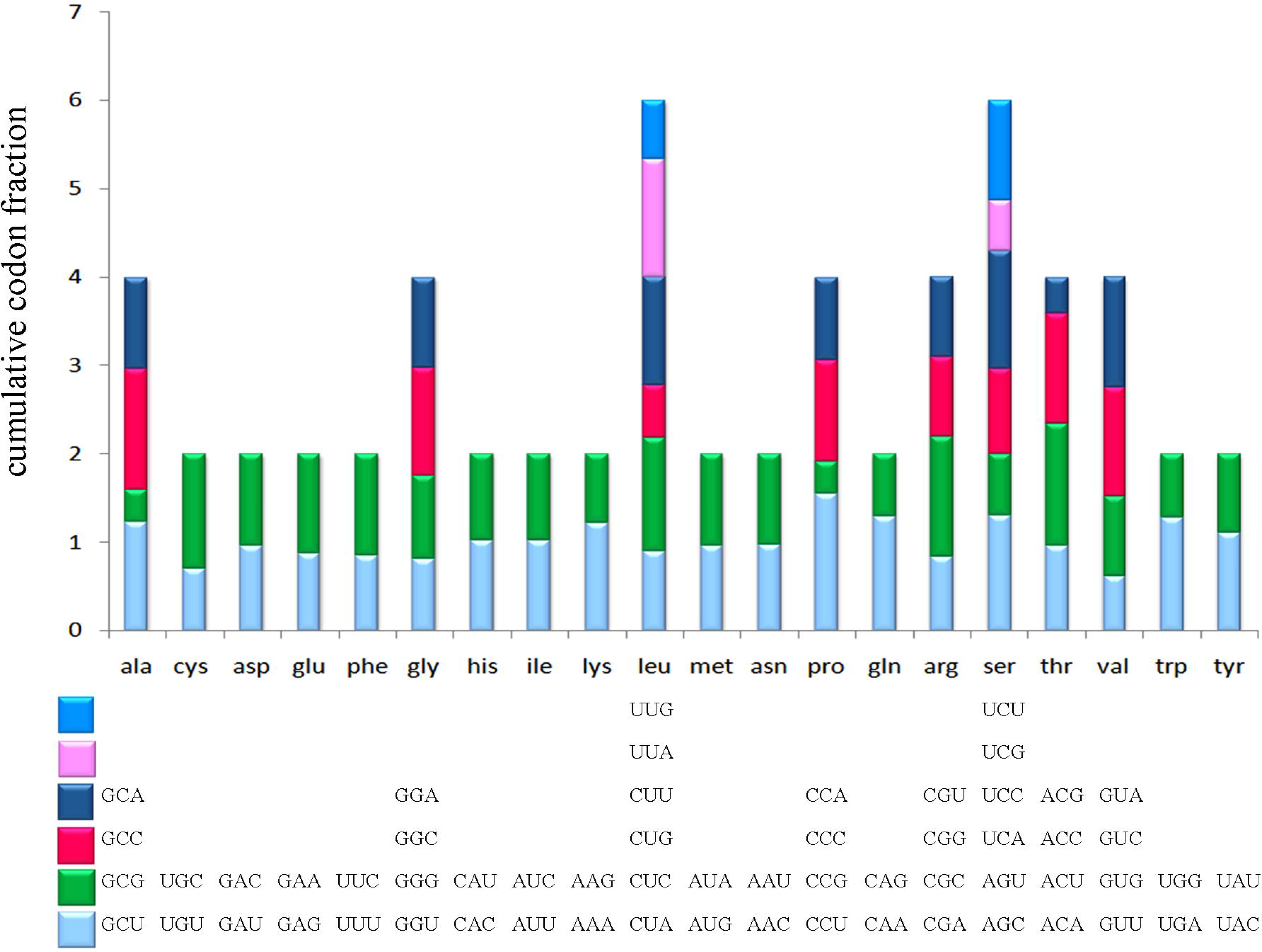
(a) ENc vs GC3 plot revealed the presence of translational selection pressure on the PCGs of select Leiothrichidae family mitogenomes, (b) RSCU analysis of *Turdoides affinis* mitogenome. X-axis represents the codon families with different colour patches. Cumulative codon fraction isplotted on Y-axis.

Moreover, values of WWC, SSU, WWU and SSC calculated from the RSCU tables for detecting the translational efficiency clearly indicated the preference of WWC and SSC over WWU and SSU. This pointed towards the selection of C between the pyrimidines (C or U) at the third position of codons. Calculated P2 values were greater than 0.5 for all the protein-coding genes in the investigated Leiothrichidae mitogenomes (Table 1, Supplementary file 3) signifying the pivotal role of translational efficiency in dictating the codon usage pattern. Translational selection along with translational efficiency plays a pivotal role in natural selection escorting towards codon preference^35^. Inclination towards translational efficiency also leads to favor codons matching with the restricted anticodon repertoire of mitochondrial tRNAs^41^ enhancing their competence in the last phase of the central dogma. The nucleotide at the third degenerating position of the codon is responsible for the superlative codon-anticodon binding energy^38^. Previous studies have found that, U is preferred at the third position specifically when G or C is present in the first two positions. On the contrary, when the first two positions are taken by A or U; C is the ‘right choice’ (at third position)^38^. Thus, translational efficiency can be characterized by the P2 index, which allows us to choose between the pyrimidines in codons with UU, UA, AA, AU, GG, GC, CG and CC in the first and second position. Results from this analysis clearly (p2>0.5) validated the accountability of translational efficiency suggesting that those genes may have gone through several adaptations with rapid changes in their expression level. The cumulative effect of all these has enhanced the capability of mitochondrial metabolic processes substantiating boisterous nature of these birds. This result showed similarity with a previous study on Dragonflies where the increased mitochondrial capacity aided their elevated flight ability^42^.

### Comparative mitochondrial genomics

The select mitogenomes were compared with *T. affinis* mitogenome (*do novo* assembled)for tRNA anticodons, start and stop codons, strand variability, intergenic and overlapping regions, GC/AT skew and RSCU (Supplementary file 3). The comparative anticodon analysis revealed an identical pattern of anticodon usages for all tRNAs among the investigated mitogenomes. Most of the protein-coding genes start with ATG in all the mitogenomes. AGG, AGA, TAG and TAA were identified as stop codons for most of the PCGs; however, in every mitogenome, there were some stop codons, which could not be perfectly identified (Supplementary file 3). Strand variation property showed an exact pattern among the studied mitogenomes Both rRNAs were on positive (+) strand. Except for nad6, all other PCGs were also on the positive (+) strand. While tRNA Q, tRNA N, tRNA C, tRNA Y, tRNA A, tRNA S2, tRNA P and tRNA E were located on the negative (–) strand, the other tRNAs were on positive (+) strand. RSCU based analysis revealed GCC(A), UGC(C), UUC(F), GGA(G), AAA(K), CUA(L), UUA(L), AUA(M), CCU(P), CAA (Q), AGC(S), UCC(S), ACC(T) and GUA(V) as optimal codons in the investigated mitogenomes. The comparative analysis of intergenic and overlapping regions also revealed identical pattern among studied mitogenomes. Intergenic regions were found between trnL2(taa) and nad1, nad1 and trnI(gat), trnI(gat) and trnQ(ttg), nad2 and trnW(tca), trnA(tgc) and trnN(gtt), trnS2(tga) and trnD(gtc), trnD(gtc) and cox2, atp6 and cox3, nad3 and trnR(tcg), nad4 and trnH(gtg), trnL1(tag) and nad5, nad5 and cob, cob and trnT(tgt), trnT(tgt) and trnP(tgg) (CR1) along with trnP(tgg) and nad6 (Supplementary file 3). The intergenic region between trnT(tgt) and trnP(tgg) were longest ranging from 939 bp (*G. cineraceus*) to 1139bp (*G. albogularis*). Overlapping regions were much shorter, ranging from 2 to 5 bp (Supplementary file 3). The highest overlapping length was between nad4l and nad4 (4bp for *G. poecilorhynchus* voucher B33 and *T. affinis,* and 5 bp for other mitogenomes(Supplementary file 3). These characters revealed identical patterns for the aforementioned characteristics in the compared mitogenomes.

### Comparative tRNA structure analysis of *T. affinis de-novo* mitogenome

The wobble base pairing which does not follow the Watson-Crick base pairing rule is of immense importance in studying the tRNA structure often substituting GC or AT base pairs contributing to thermodynamic stability^43^. All these features together affect several biological processes^44^. Studies have reported that, RNA binding proteins generally adhere to G-U sites differing from the Watson-Crick or other mis-matched base-pair pattern^45^. Hence, while understanding the exact functional features of mitogenomes, tRNA acts as a pivotal tool ^45, 46^. In *T. affinis*, all the tRNAs were folded into classic secondary clover-leaf structure. In *T. affinis*mitogenome, though Watson-Crick base pair dominated, wobble base pairs were also detected (Fig. 7). For instance, trnA, trnC, trn E, trnN, trnP, trnS1, trnS2 and trnY had wobble base pairing at the acceptor arm. Moreover, trnA, trnC, trnS1, trnQ, trnP and trnN contained wobble base pairing at TΨC stem. Same G-U base pairs were also detected at the anticodon stem of trnC, trnP, trnQ, trnS1 and trnT. Three consecutive wobble base pairs were detected at DHU loop of trnS2. Along with, some other mismatched base pairs were also found in trnD, trnE, trnG, trnH, trnM and trnL2. Among other species considered in the present study, all the tRNAs were found to be folded in clover-leaf structure with dominating Watson-Crick base pairing.

**Figure 7.**
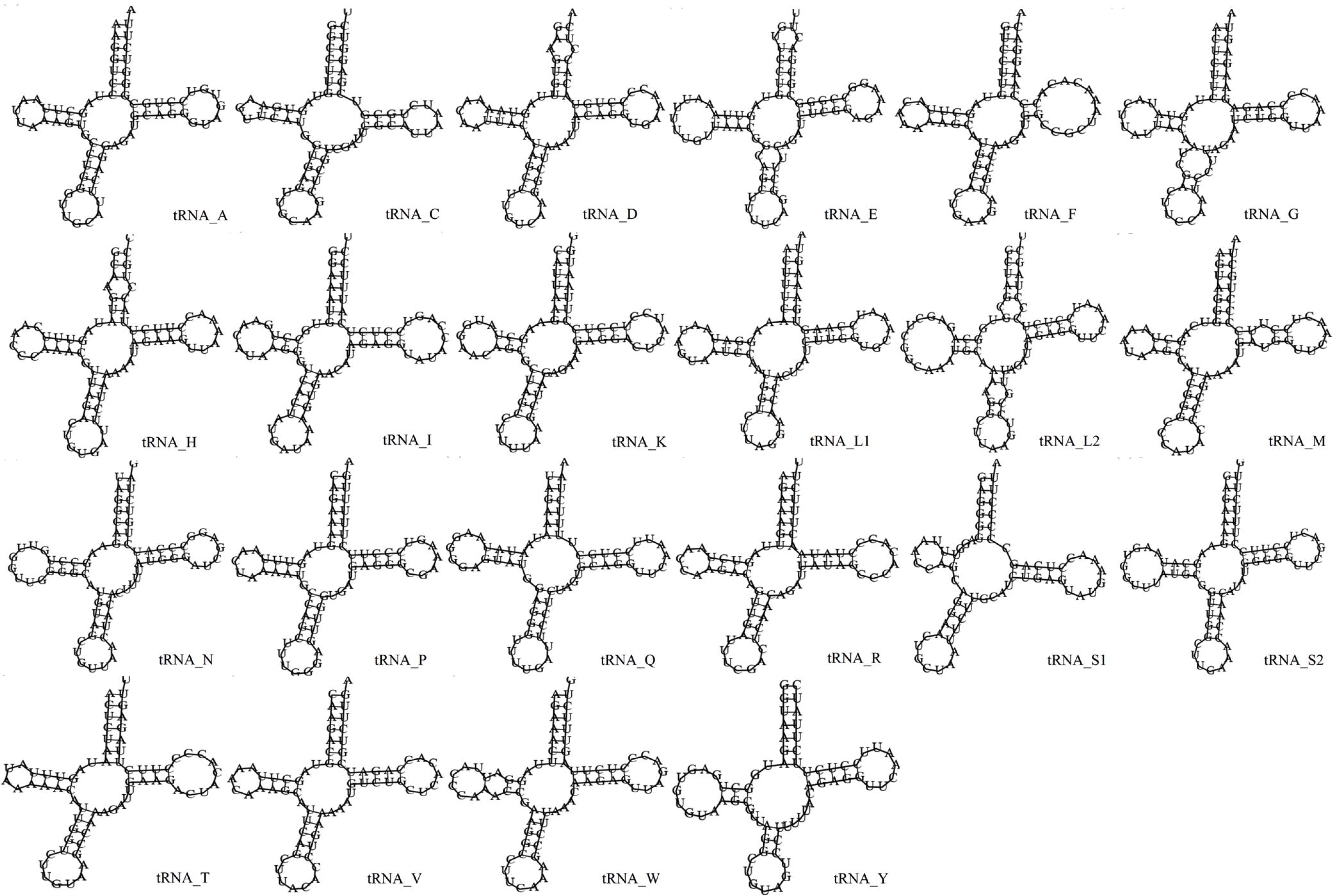
Putative secondary structures of *Turdoides affinis* tRNAs. tRNAs are specified with respective single letter amino acid codes.

### Control region of *T. affinis de-novo* mitogenome

Vertebrate mitochondrial Control Region (CR) is divided in to three domains (I, II and III)^46^. Domain I contains three Extended Terminal Associated Sites (ETAS) proceeded by a C-rich region. Domain II contains F-box, E-box, D-box, C-box and B-box^47^. For birds, there is a special Bird Similarity Box (BSB) present in the left side of B-box in domain II. Domain III consists of three conserved sequence blocks (CSB-I, II, III). Domain III also contains replication origin of H-strand along with bi-directional promoters for both H- and L-strand transcription. Domain II is supposed to be more conserved than Domain I and III^47^. Among all the investigated species of Leiothrichidae family, a duplication of CR was observed. All the aforementioned domains and conserved boxes were identified through sequence alignment. Conserved BSB was prominent in both CR1 and CR2 (Fig. 8). No tandem repeats were found among the CRs. Duplication of CR region is important in regulation of replication and transcription within mitochondrial genome^46^. Moreover, duplicated CR is also associated with extended longevity of bird species^48^. Thus, the present study reported the genetic features of duplicated CR among select Leiothrichidae members including *T. affinis,* which will further be helpful in evolutionary analysis of this group.

**Figure 8.**
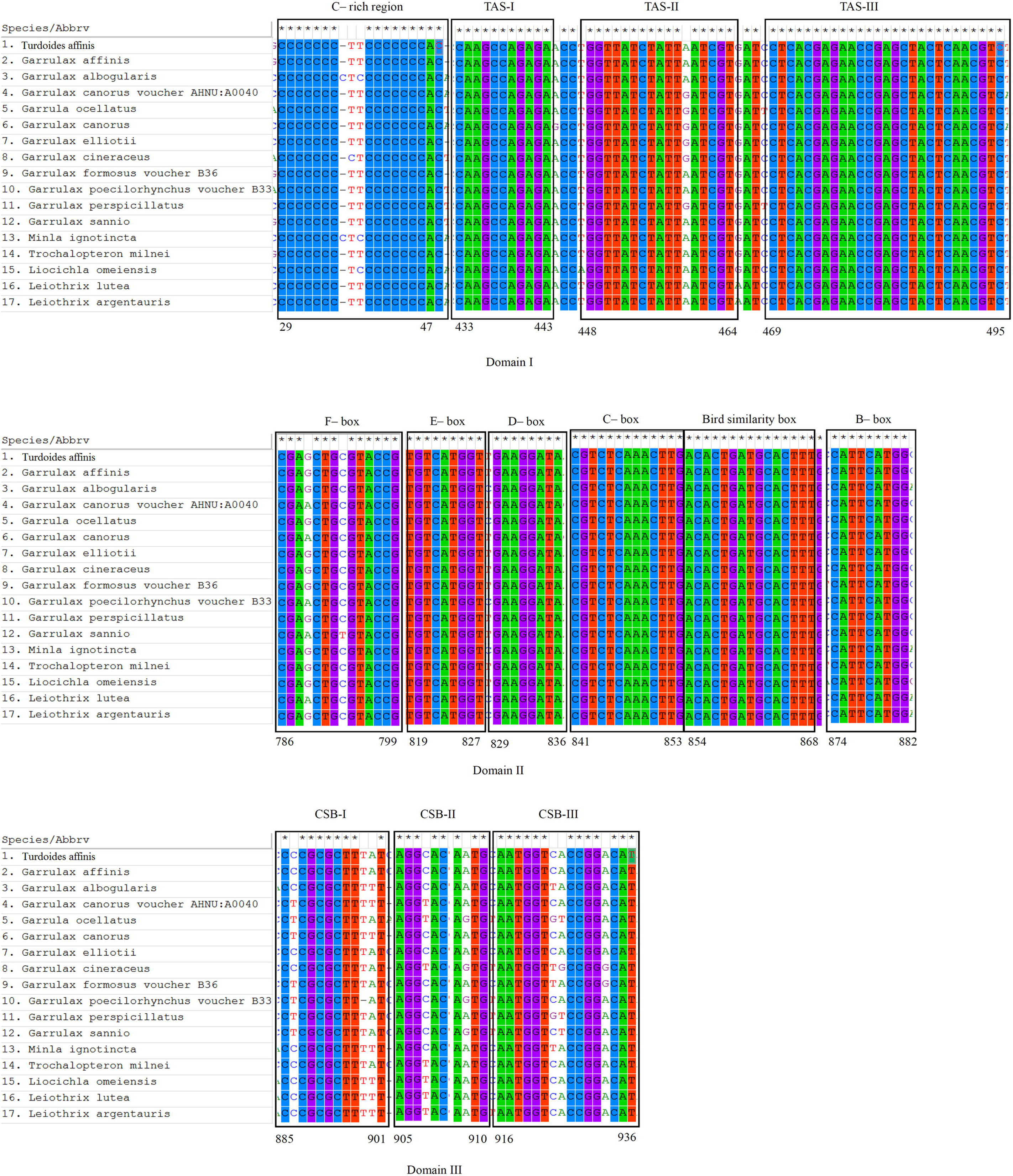
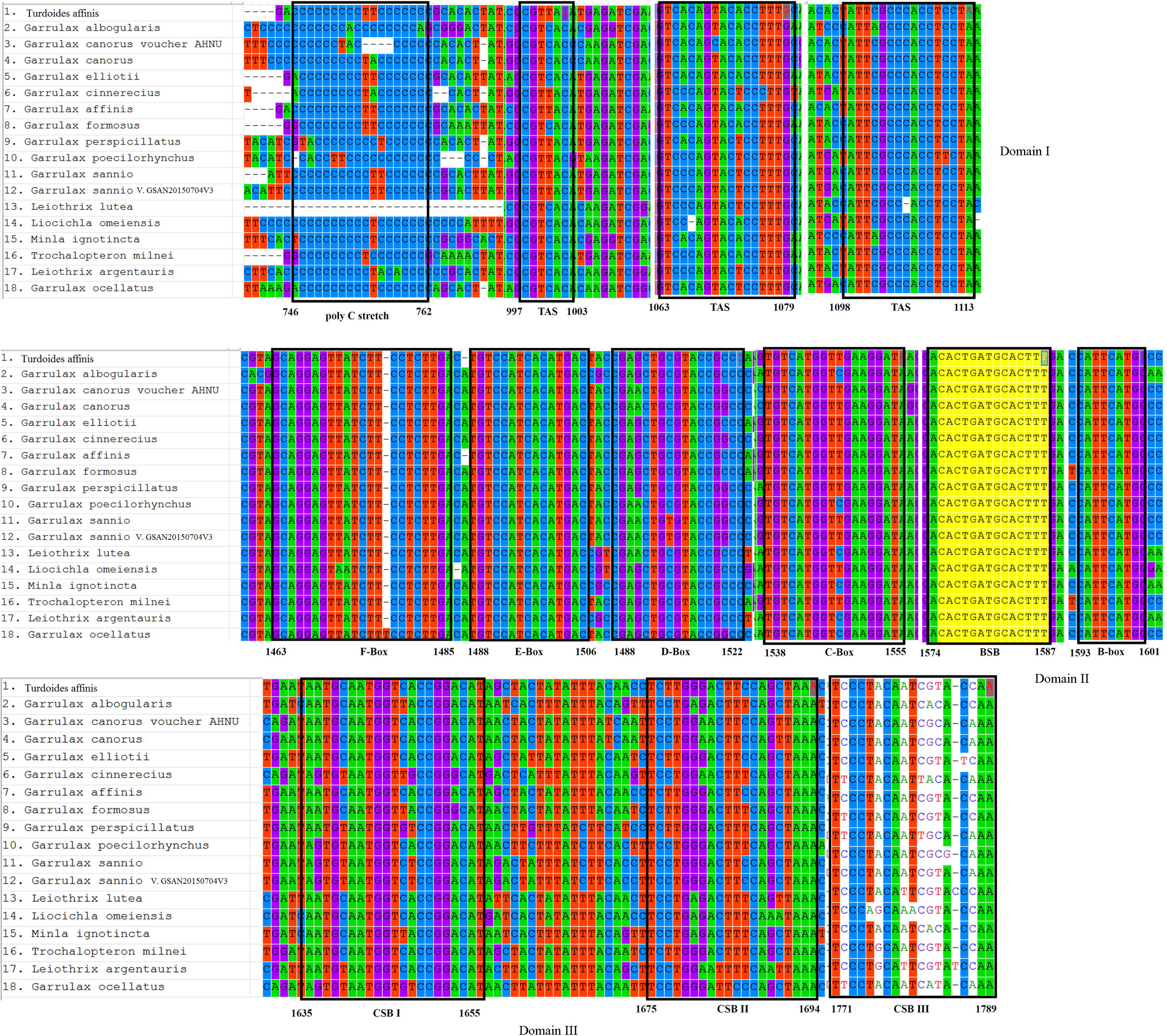
Domains and boxes identified in (a) CR1 and (b) CR2 of select Leiothrichidae family mitogenomes. Regions of the identified boxes are given at the lower part of each box.

### Evolutionary analysis

Genetic and evolutionary distance calculation revealed higher K2P distance in nad2, atp8 and nad6 whereas cox1 possessed the lowest value among the Leiothrichidae family (Fig. 9). Regarding the ka/ks analysis, the average synonymous substitution rate (Ks) for nd2 gene was highest whereas non-synonymous substitution rate (ka) was highest for atp8. ⍰(ka/ks) values of the protein-coding genes ranged from 0.014 to 0.183 and was in the following order- cox1<cox3<nad1<cob<nad4l<atp6<nad4<cox2<nad3<nad5<nad6<nad2<atp8 (Supplementary file 3). Lowest K2P distance for cox1 gene indicated towards its conserved nature among the Leiothrichidae family. Moreover, this also implicated the preference of NADH:ubiquinoneoxidoreductase (complex I) over the succinate dehydrogenase complex in electron transport chain all through the examined family. Complex I is responsible for exporting four H+ ion out of the mitochondrial matrix participating in the generation of H+ gradient across the mitochondrial membrane, which ultimately speeds up ATP generation whereas this mechanism is totally absent when complex II is used^49^. Moreover, the non-synonymous substitution rate of cox1 was also found to be least, indicating its conserved nature in mitochondrial machinery of Leiothrichidae family. Mitochondria associated NADH dehydrogenase 2 encoded by the nad2 gene is also a subunit of complex 1 located in the inner mitochondrial membrane and is the largest complex of ETS^49^. The high synonymous substitution rate (Ks) of nad2 gene further indicated the preserved amino acid component of complex I. Highest Ka value of atp8 pointed to a highly variable nature of this protein indicating the erratic nature of mitochondrial atp8 among vertebrates^50^. The ⍰ (ka/ks) values of all the protein-coding genes were <1 suggesting the persistence of purifying selection against deleterious mutation. Thus, the evolutionary analysis aided in understanding the influence of natural selection manipulating species evolution along with the interaction between selection and mutational pressure responsible for protein evolution which has been already suggested^39^.

**Figure 9.**
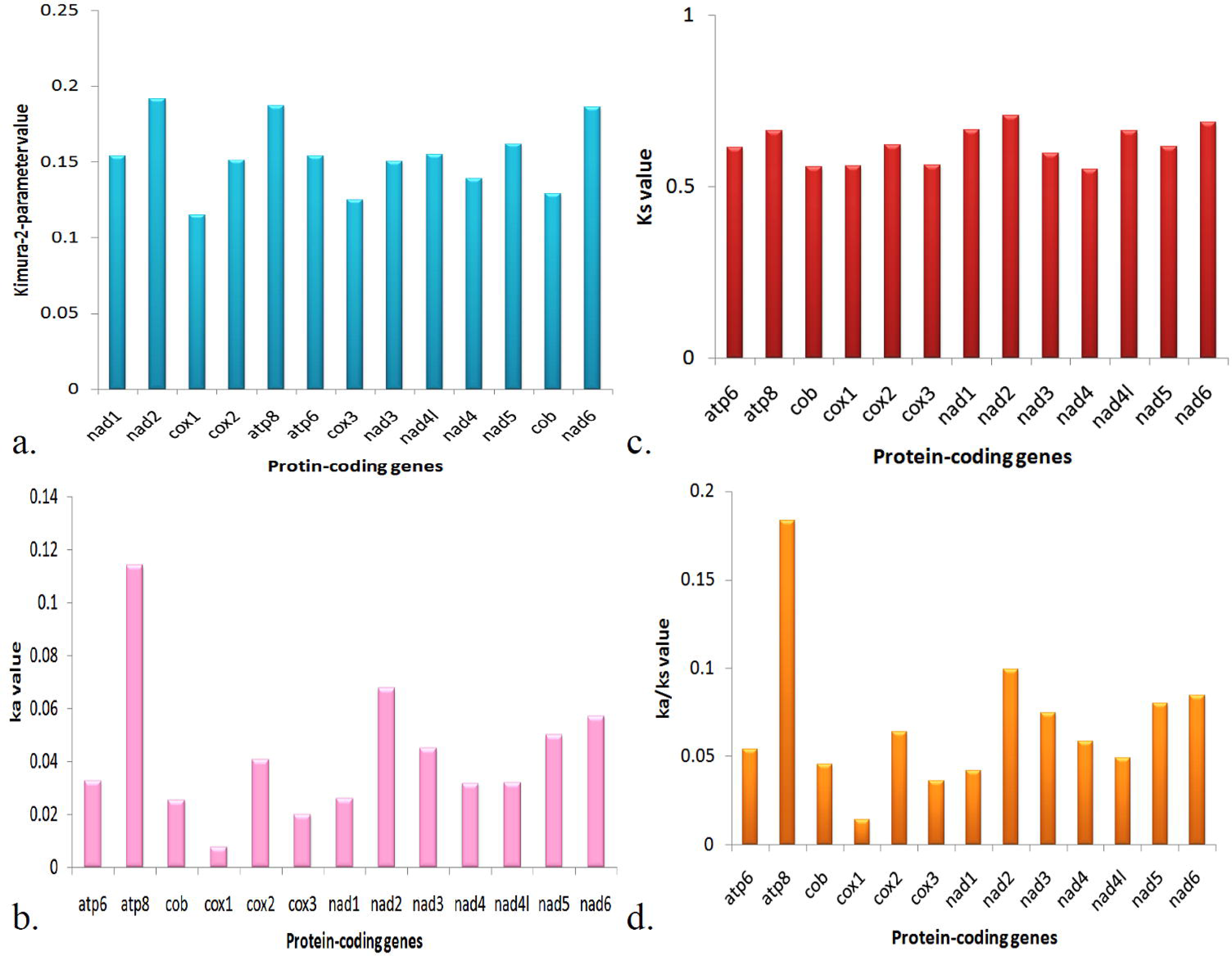
Genetic and evolutionary distance among the PCGs of select species of the Leiothrichidae family. (a) K2P distance calculation, (b)Ks values, (c)Ka values and (d)Ka/Ks values of mitochondrial PCGs among investigated species of Leiothrichidae family.

## Conclusion

We report for the first time the complete mitogenome of *Turdoides affinis*(MN848144).We find the *de-novo* assembly approach more appropriate than a reference-based assembly approach. The comparative mitogenomics of the Leiothrichidae family reveals their preference towards AT-rich codons as well as the persistence of translational efficiency. tRNA analysis shows the dominance of Watson-Crick base pairing with a few exceptions of wobble base pairing. Duplicated control regions are found among Leiothrichidae family mitogenomes that may help in their extended longevity. Evolutionary analysis confirms thatprotein-coding genes are under purifying selection pressure. Genetic distance and variation analysis indicate the dominance of NADH dehydrogenase complex-I in the electron transport system of *T. affinis*.Our limited phylogenetics results suggest that *T. Affinis* is closer to *Garrulax*.

## Supporting information

Supplementary file 1. Description of complete mitogenome of Turdoides affinis assembled through reference based method.

Supplementary file 2. Phylogenetic tree with complete mitogenome of Turdoides affinis (reference based assembly).

Supplementary file 3. Comparative mitogenomics among compared Leiothrichidae family mitogenomes.

## Acknowledgments

We thank the Ministry of Environment, Forest and Climate Change, Govt. of India for financial support. We are also thankful to the Tamil Nadu Forest Department for providing the permissions and support to conduct the study.

## Author Contribution

RPS collected samples and conceived the idea. RPS, SKS, PD and SDR designed the experiments and generated DNA data. IS analysed the data. RPS and IS wrote the manuscript and generated all the figures. All authors reviewed the manuscript.

---

**Supplementary file 1**. Description of complete mitogenome of *Turdoides affinis* assembled through reference based method.

**Supplementary file 2.** Phylogenetic tree with complete mitogenome of *Turdoides affinis*

(reference based assembly).

**Supplementary file 3.** Comparative mitogenomics among compared Leiothrichidae family mitogenomes.

