## Supplementary file 1. Description of complete mitogenome of Turdoides affinis assembled through reference based method. for "Comprehensive bioinformatic analysis of newly sequenced *Turdoides affinis* mitogenome reveals the persistence of translational efficiency and dominance of NADH dehydrogenase complex-I in electron transport system over Leiothrichidae family"

**Description of complete mitochondrial genome of *Turdoides affinis* assembled using a reference genome of *Leiothrix argentauris***

The raw paired end data generated from Nextseq 550 was analysed by bcl2fastq 2.0+ software to trim the over-represented adapter sequences before further processing. This software cleaned the adapter sequences and converted bcl files into FastQ format. FastQC (https://www.bioinformatics.babraham.ac.uk/projects/fastqc/) was used to obtain the quality control report of the sequences. The mitochondrial genome of another Leiothrichidae family *Leiothrix argentauris* was used as a reference genome for aligning our sequence in BWA (<http://bio-bwa.sourceforge.net/>). This aligned data in bam format was used in samtools (http://www.htslib.org/) for indexing step. Mapped short reads were viewed in Integrative Genomics Viewer (http://software.broadinstitute.org/software/igv/). The aligned regions were extracted through samtools which gave a consensus sequence for complete mitochondrial genome of *Turdoides affinis*. Annotation of the obtained mitogenome was performed by MITOS using genetic code 2 (for vertebrates). Protein coding genes (PCGs), rRNA as well as tRNAs were annotated at this step. tRNAscan-SE v1.3.1 (Lowe and Eddy, 1997) was used to predict the tRNA present within the genome and the obtained result was compared with MITOS which gave a perfect match and further validated the annotation strategy of MITOS.

The complete mitochondrial genome of *Turdoides affinis* was sequenced and analyzed (Fig. SF1_A). The total size of the mitogenome was 16,861 bp with 47% GC composition (53% AT). AT and GC skew of the sequenced mitogenome was found to be 0.05 and -0.14 respectively. Total thirteen protein coding genes were detected along with two rRNA genes (small subunit ribosomal rrns and large subunit ribosomal rrnl). Twenty-two tRNA genes were detected encoding 20 standard amino acids (two tRNAs were there for serine and leucine). All the protein coding genes were on positive strand (+) strand except nad6. trnQ, trnA, trnN, trnC, trnY, trnS2, trnP and trnE were found to be located on negative (-) strand whereas rest of the tRNAs were on positive strand.

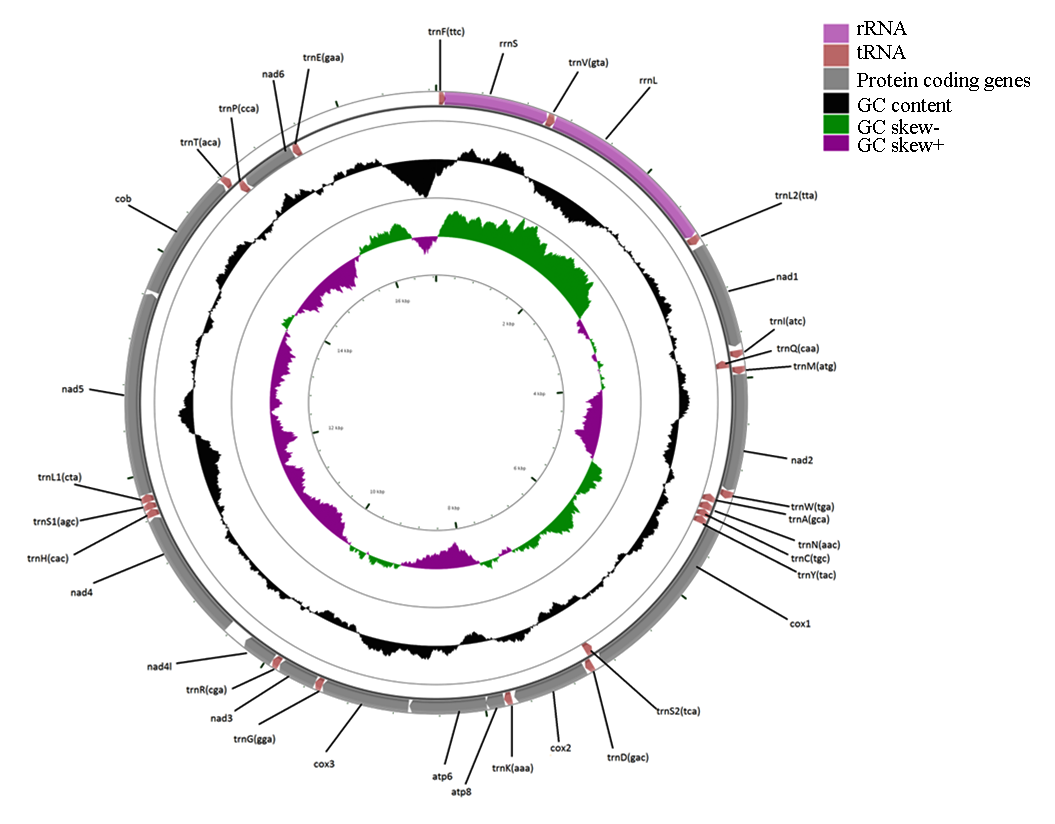

SF1_A: Circular genome view of complete mitogenome of *Turdoides affinis* obtained using reference based assembly.

Genes in *Turdoides affinis* was found arranged in the following manner which was quite identical among Leiothrichidae family: trnF-rrnS-trnV-rrnL-trnL2-nad1-trnI-trnQ-trnM-nad2-trnW-trnA-trnN-trnC-trnY-cox1-trnS2-trnD-cox2-trnK-atp8-atp6-cox3-trnG-nad3-trnR-nad4l-nad4-trnH-trnS1-trnL1-nad5-cob-trnT-trnP-nad6-trnE.

Codon usage analysis indicated UUC (Phe), CUC and CUA (Leu), AUC (Ile), GUC and GUA (Val), UAC(Tyr), CAC (His), CAA(Gln), AAC (Asn), AAA (Lys), GAC (Arg), GAA (Glu), UCA and UCC(Ser), CCA, CCC and CCU(Pro), ACC and ACA (Thr), GCC and GCA (Ala), UGC(Cys), CGC(Arg), UCC and UCA(Ser) and GGA and GGC (Gly) were frequently used codons.

Four overlapping regions were found varied in length from 1bp to 4 bp (Table SF1_B). Largest overlapping regions were between nad1 and trnI (atc) as well as atp8 and atp6. The intergeneric spacer sequences spread over 25 regions and their length varied from 1bp to 825bp. The largest spacer sequence (length 825 bp) was found between trnT(aca) and trnP(cca). Similar analysis on the other considered mitogenomes revealed an identical pattern of intergenic and overlapping regions. The length of the intergenic spacer region between trnT and trnP were highest among all the examined mitogenomes rather than other regions. The length of that region varied from 1139 bp for *G. albogularis* (maximum) to 825 bp in *Turdoides affinis.*

Table SF1_B: Annotation of *Turdoides affinis* complete mitogenome assembled using reference genome.

| **Locus name** | **Start Position** | **End position** | **Strand** | **Intergenic Nucleotides** | **Size (bp)** | **Anti-codon** | **WC**  **Codon** | **Initiation codon** | **Terminating Codon** |
| --- | --- | --- | --- | --- | --- | --- | --- | --- | --- |
| trnF(ttc) | 5 | 69 | + | 1 | 65 | GAA | UUC | - | - |
| rrnS | 71 | 1048 | + | -1 | 978 | - | - | - | - |
| trnV(gta) | 1048 | 1118 | + | 0 | 71 | UAC | GUA | - | - |
| rrnL | 1119 | 2718 | + | 1 | 1600 | - | - | - | - |
| trnL2(tta) | 2720 | 2793 | + | 33 | 74 | UAA | UUA | - | - |
| nad1 | 2827 | 3762 | + | -4 | 936 | - | - | ATC | AGA |
| trnI(atc) | 3759 | 3830 | + | 5 | 72 | UCG | AUC | - | - |
| trnQ(caa) | 3836 | 3906 | - | -1 | 71 | UUG | CAA | - | - |
| trnM(atg) | 3906 | 3974 | + | 0 | 69 | CAU | AUG | - | - |
| nad2 | 3975 | 5009 | + | 4 | 1035 | - | - | ATG | TAG |
| trnW(tga) | 5014 | 5084 | + | 1 | 71 | UCA | UGA | - | - |
| trnA(gca) | 5086 | 5154 | - | 8 | 69 | UGC | GCA | - | - |
| trnN(aac) | 5163 | 5235 | - | 1 | 73 | GUU | AAC | - | - |
| trnC(tgc) | 5237 | 5302 | - | 0 | 66 | GCA | UGC | - | - |
| trnY(tac) | 5303 | 5372 | - | 1 | 70 | GUA | UAC | - | - |
| cox1 | 5374 | 6912 | + | 0 | 1539 | - | - | ATA | AGG |
| trnS2(tca) | 6913 | 6987 | - | 5 | 75 | UGA | UCA | - | - |
| trnD(gac) | 6993 | 7061 | + | 10 | 69 | GUC | GAC | - | - |
| cox2 | 7072 | 7740 | + | 26 | 669 | - | - | ATG | TAA |
| trnK(aaa) | 7767 | 7834 | + | 1 | 68 | UUU | AAA |  |  |
| atp8 | 7836 | 7997 | + | -4 | 162 | - | - | ATC | TAA |
| atp6 | 7994 | 8674 | + | 8 | 681 | - | - | ATG | AGA |
| cox3 | 8683 | 9465 | + | 1 | 783 | - | GGA | - | - |
| trnG(gga) | 9467 | 9535 | + | 0 | 69 | UCC | - | - | - |
| nad3 | 9536 | 9883 | + | 4 | 348 | UCG | - | ATT | TAA |
| trnR(cga) | 9888 | 9957 | + | 9 | 70 | - | CGA |  |  |
| nad4l | 9967 | 10248 | + | 6 | 282 | - | - | ATC | AGA |
| nad4 | 10255 | 11613 | + | 9 | 1359 | - | - | ATC | TAA |
| trnH(cac) | 11623 | 11691 | + | 0 | 69 | GUG | CAC | - | - |
| trnS1(agc) | 11692 | 11757 | + | 0 | 66 | GCU | AGC | - | - |
| trnL1(cta) | 11758 | 11828 | + | 0 | 71 | UAG | CUA | - | - |
| nad5 | 11829 | 13634 | + | 20 | 1806 | - | - | ATA | TAG |
| cob | 13655 | 14788 | + | 10 | 1134 | - | - | ATC | TAA |
| trnT(aca) | 14799 | 14867 | + | 825 | 69 | UGU | ACA | - | - |
| trnP(cca) | 15693 | 15758 | - | 9 | 66 | UGG | CCA | - | - |
| nad6 | 15768 | 16280 | - | 5 | 513 | - | - | ATC | TAG |
| trnE(gaa) | 16286 | 16357 | - | 0 | 72 | UUC | GGA | - | - |
| Control region | 16358 | 17882 | - | - | 1529 | - | - | - | - |
|  | 1 | 4 |  | - |  | - | - | - | - |

**Control region**

The control region (CR) was 1529 bp long (Table SF1_B ) with 55.9% of AT composition. This AT rich CR was also found among other studied Leiothrichidae mitogenomes (Fig SF1_C).
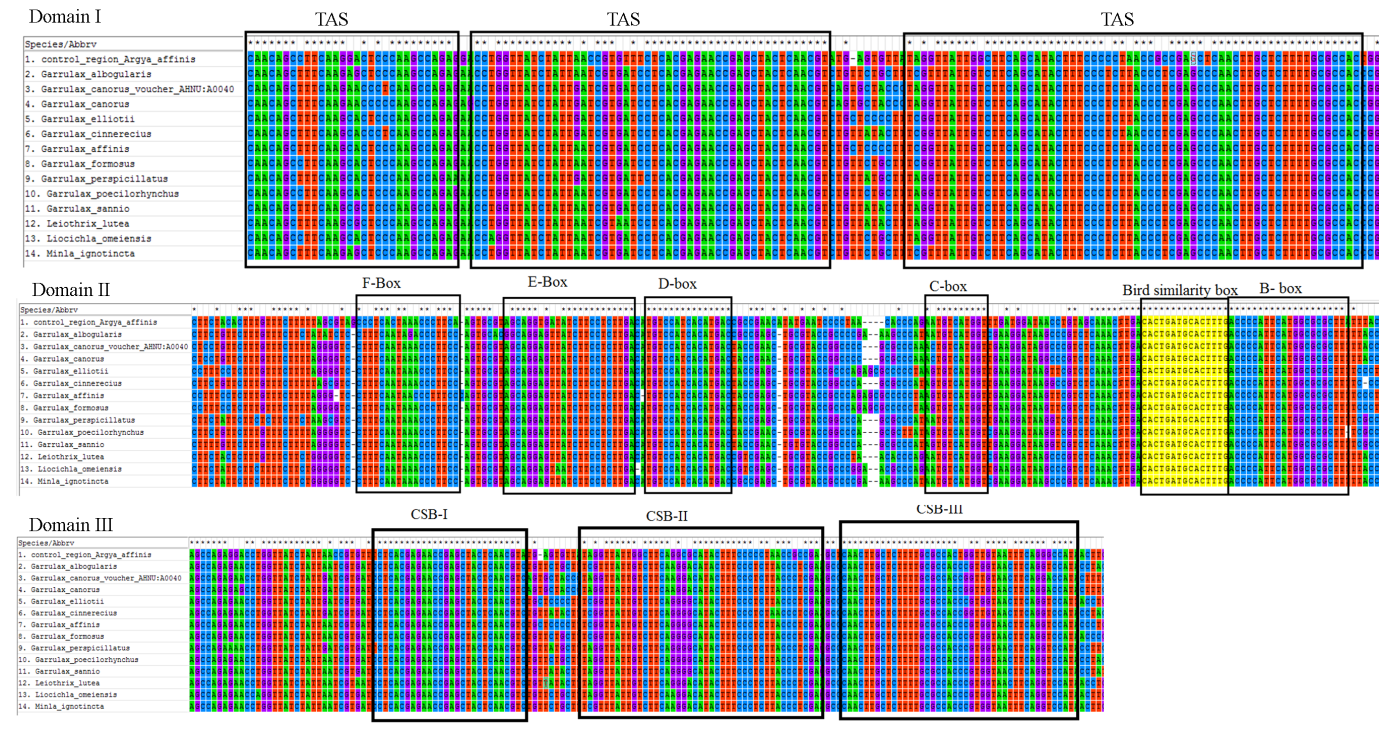

Fig. SF1_C: Control region comparison of mitogenomes from considered species

The length of the CR of *Turdoides affinis* was 1529 bp with 53.86% of AT. Three conserved parts were found in the domain I. Domain II contained highly conserved D box, B-box and bird similarity box. However, F-box, E-box and C-box were not that much conserved and were found to contain variable nucleotide sites. Three CSB (I, II, III) were also found in the Domain III region however, they also contained a few variable nucleotide site (SF1_C).

**Variations in the tRNA secondary structures**

The tRNA secondary structures from all the investigated mitogenomes were compared to explore variations among different species. Most interestingly, the secondary structures of tRNA F (phenylalanine) and tRNA H (histidine) in *Turdoides affinis* were totally different from others compared species. Besides, tRNA A (alanine), tRNA I (isoleucine) and tRNA S1 (serine) also showed considerable variations from others compared species.

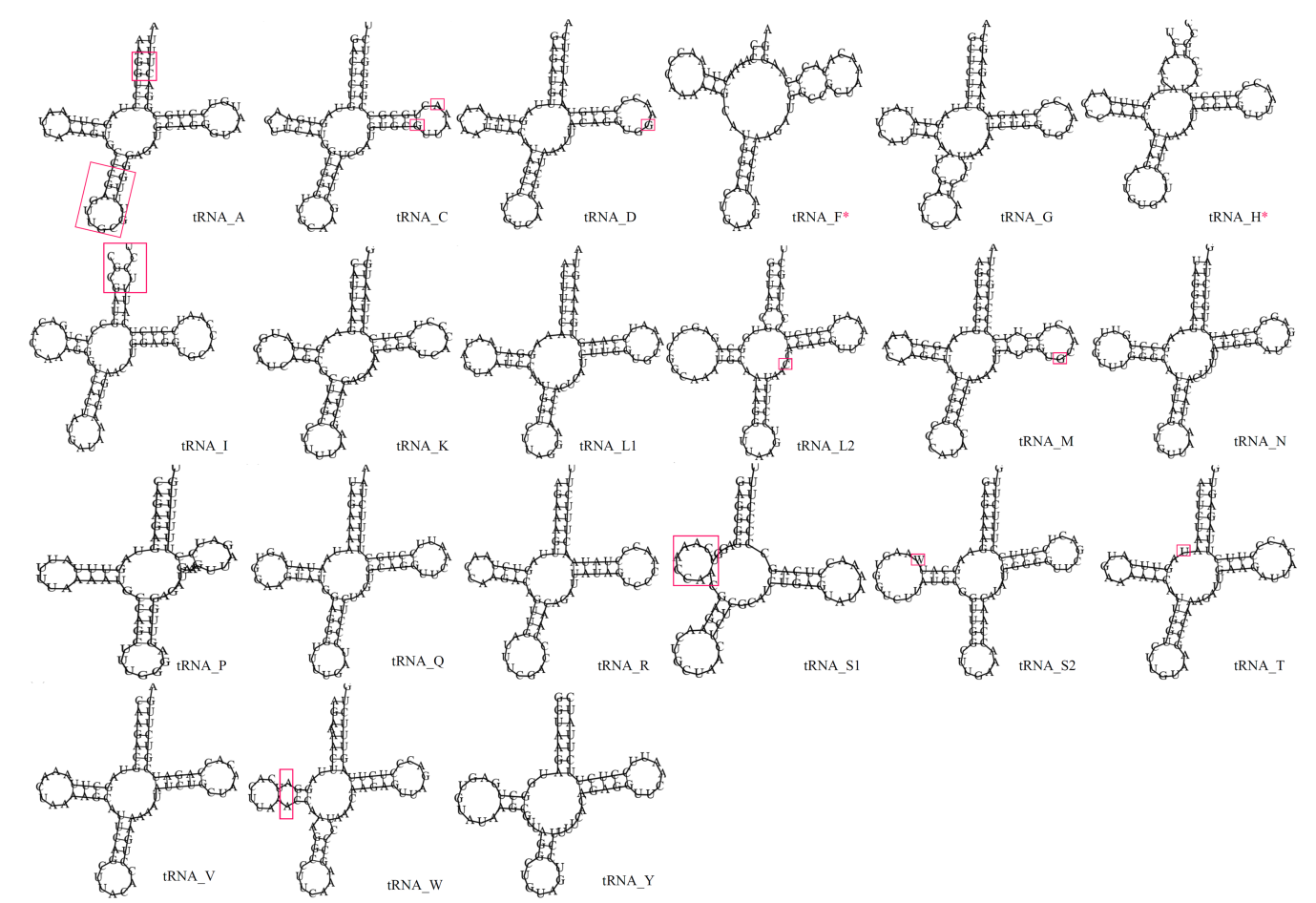

SF1_D: tRNA secondary structures from *Turdoides affinis* complete mitogenome assembled through reference based approach.

Some tRNAs varied for only one nucleotide whereas, rest showed similarities with at least one another species. Most sites of each tRNA showed normal Watson crick base pairing however, wobble base pairing was also observed in the anti-codon stem and TΨC stem of tRNAs. Conserved secondary structure for tRNA G, tRNA K, tRNA L1, tRNA N, tRNA P, tRNA Q, tRNA R, tRNA V and tRNA Y was found among all studied mitogenomes.
