## Supplementary file 2. Phylogenetic tree with complete mitogenome of Turdoides affinis (reference based assembly). for "Comprehensive bioinformatic analysis of newly sequenced *Turdoides affinis* mitogenome reveals the persistence of translational efficiency and dominance of NADH dehydrogenase complex-I in electron transport system over Leiothrichidae family"

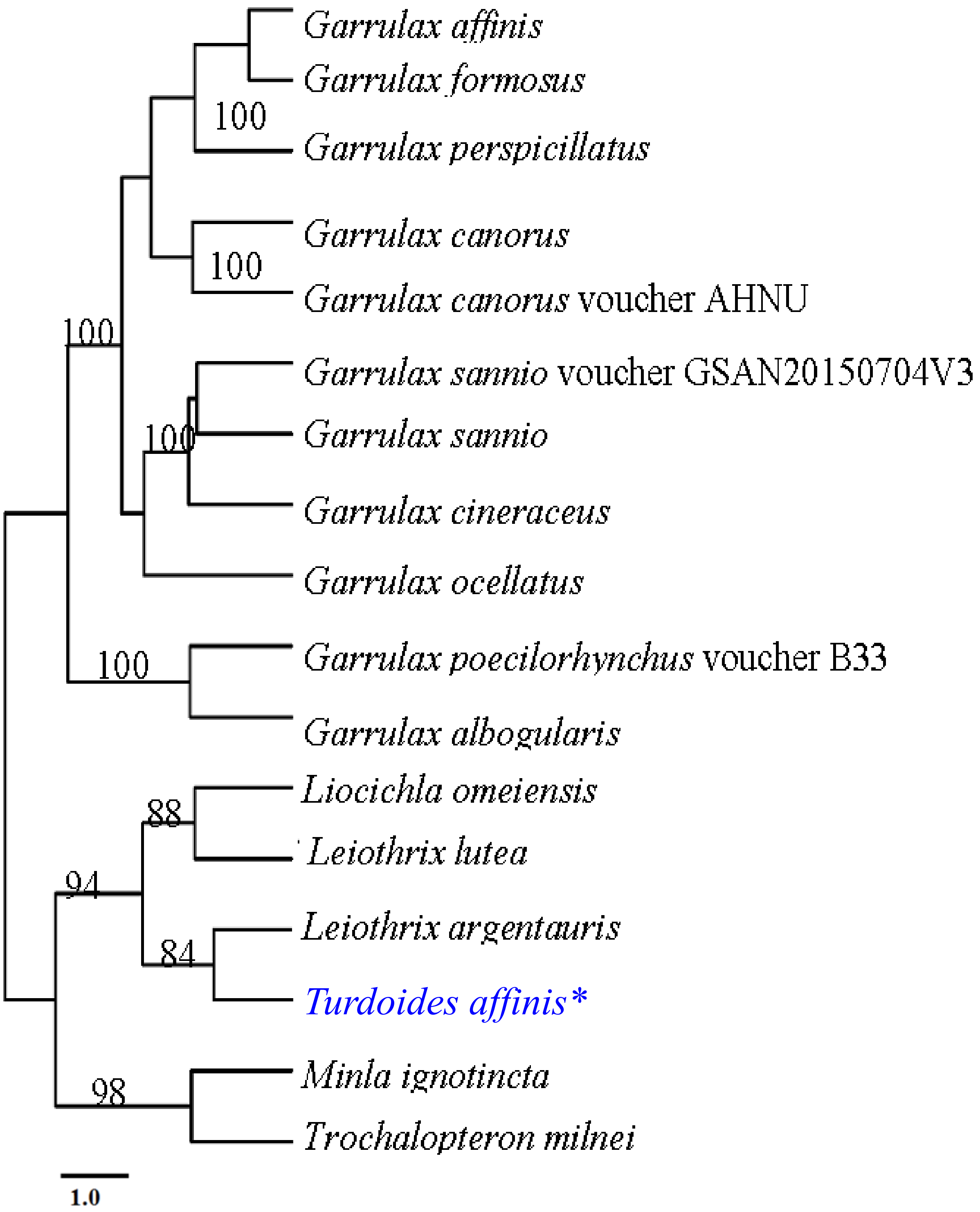


Supplementary File 2: Phylogenetic tree based on reference based assembled complete mitogenome of *T. affinis.* Maximum likelihood method with 1000 bootstrap value was used to build the tree.
